# Restrictor slows early transcription elongation to render RNA polymerase II susceptible to termination at non-coding RNA loci

**DOI:** 10.1101/2025.01.08.631787

**Authors:** Claudia A. Mimoso, Hanneke Vlaming, Nathalie P. de Wagenaar, Karen Adelman

**Affiliations:** Department of Biological Chemistry and Molecular Pharmacology, Harvard Medical School, Boston, MA 02115, USA; Department of Genetics, Yale School of Medicine, New Haven, CT USA; Division of Genome Biology & Epigenetics, Institute of Biodynamics and Biocomplexity, Utrecht University, Utrecht, The Netherlands

**Author notes:** Co-first authors (Claudia A. Mimoso, Hanneke Vlaming).

## Abstract

The eukaryotic genome is broadly transcribed by RNA polymerase II (RNAPII) to produce protein-coding messenger RNAs (mRNAs) and a repertoire of non-coding RNAs (ncRNAs). Whereas RNAPII is very processive during mRNA transcription, it terminates rapidly during synthesis of many ncRNAs, particularly those that arise opportunistically from accessible chromatin at gene promoters or enhancers. The divergent fates of mRNA versus ncRNA species raise many questions about how RNAPII and associated machineries discriminate functional from spurious transcription. The Restrictor complex, comprised of the RNA binding protein ZC3H4 and RNAPII-interacting protein WDR82, has been implicated in restraining the expression of ncRNAs. However, the determinants of Restrictor targeting and the mechanism of transcription suppression remain unclear. Here, we investigate Restrictor using unbiased sequence screens, and rapid protein degradation followed by nascent RNA sequencing. We find that Restrictor promiscuously suppresses early elongation by RNAPII, but this activity is blocked at most mRNAs by the presence of a 5’ splice site. Consequently, Restrictor is a critical determinant of transcription directionality at divergent promoters and prevents transcriptional interference. Finally, our data indicate that rather than directly terminating RNAPII, Restrictor acts by reducing the rate of transcription elongation, rendering RNAPII susceptible to early termination by other machineries.

## INTRODUCTION

The collection of mRNA molecules present in a cell dictates protein production and thus cellular behavior and function. Establishing the appropriate repertoire of mRNAs, and preventing the formation and accumulation of spurious transcripts, involves precise, coordinated regulation of transcription, RNA processing and RNA decay. A central point of gene control occurs early in transcription elongation, with the establishment of promoter-proximally paused RNAPII, which is bound and stabilized by the DSIF and NELF complexes (Core and Adelman 2019; Adelman and Lis 2012). Regulated pause release, triggered by the kinase activity of P-TEFb, dissociates NELF and enables the association of a myriad of factors that facilitate RNAPII elongation across the chromatin template (Peterlin and Price 2006; Vos et al. 2018). Further, the presence of a 5’ splice site (5’ SS) within the initially transcribed sequence of most mRNAs recruits the U1 small nuclear ribonucleoprotein (snRNP), which both enhances RNAPII elongation rate and initiates the first steps in RNA splicing (Mimoso and Adelman 2023; Shine et al. 2024; Carrocci and Neugebauer 2024). Thus, at mRNA genes, cis- and trans-acting factors cooperate to prevent premature termination and stimulate the production and efficient processing of mature mRNAs (Shine et al. 2024; Bieberstein et al. 2012; Venters et al. 2019; Tudek et al. 2018; Kaida et al. 2010; Berg et al. 2012).

ncRNAs transcribed from regulatory regions in the genome, such as enhancer RNAs (eRNAs) and upstream antisense RNAs (uaRNAs, or PROMPTs), are typically much shorter and less stable than mRNAs (Preker et al. 2008; Core et al. 2008; Seila et al. 2008; Core et al. 2014; Henriques et al. 2018). Early termination within several kb of the transcription start site (TSS) is very common (Henriques et al. 2018; Lykke-Andersen et al. 2021), and many ncRNAs lack the positive sequence features (e.g., 5’ SS) and recruitment of elongation factors characteristic of mRNAs (Almada et al. 2013; Ntini et al. 2013). Indeed, preventing uncontrolled or pervasive elongation of these ncRNAs is thought to safeguard the genome by averting transcription interference and polymerase collisions that could lead to DNA damage (Cinghu et al. 2017; Flynn et al. 2016; Ogami et al. 2017). Achieving the correct balance between elongation and termination across RNAPII-transcribed loci is thus critical for appropriate gene expression and ensuring genome integrity.

Transcription termination by RNAPII is typically driven by multi-subunit protein complexes, often aided by sequence features (Shi and Manley 2015; Proudfoot 2016; Davidson et al. 2024). Well-defined termination machineries include the cleavage and polyadenylation (CPA) machinery and the Integrator complex, both of which harbor RNA endonuclease and phosphatase activities to cleave nascent RNA and remove stimulatory phosphorylation from RNAPII and elongation factors (Rodríguez-Molina et al. 2023; Wagner et al. 2023). The CPA machinery governs the formation of most canonical mRNA 3’-ends through specific recognition of polyadenylation signals (PAS) (Chan et al. 2014; Shi and Manley 2015; Proudfoot 2016). In addition, the CPA factors can act at cryptic PAS motifs that are present in ncRNAs or found within AT-rich introns, causing premature cleavage and polyadenylation (PCPA) (Kaida et al. 2010; Berg et al. 2012). Notably, PCPA often leads to the production of short polyadenylated RNAs that are selectively recognized for degradation by the exosome (Venters et al. 2019; Garland et al. 2019; Tudek et al. 2018).

In contrast, the Integrator complex has no known DNA or RNA sequence specificity, and instead broadly targets RNAPII early elongation complexes of all RNA biotypes (Lai et al. 2015; Kirstein et al. 2021; Stein et al. 2022; Lykke-Andersen et al. 2021). Structural studies reveal that Integrator interacts with DSIF, NELF and RNAPII, providing specificity for promoter-proximally paused RNAPII (Fianu et al. 2024, 2021). Integrator functions to terminate paused RNAPII that is not released by P-TEFb, ensuring that only fully elongation competent RNAPII enter gene bodies (Elrod et al. 2019; Beckedorff et al. 2020; Huang et al. 2020; Lykke-Andersen et al. 2021; Stein et al. 2022; Hu et al. 2023). In this way, Integrator attenuates the expression of select protein-coding genes and suppresses transcription of spurious ncRNAs.

The Restrictor complex has also been implicated in suppressing the expression of many ncRNAs. The central component of this complex, ZC3H4 in mammals and SU(S) in Drosophila, is a protein with two arginine rich motifs and zinc-fingers, that binds RNA with high affinity but limited specificity (Murray et al. 1997; Turnage et al. 2000). SU(S) was shown to inhibit RNA production in a manner that could be overcome by assembly of the splicing complex, although whether this involved control of transcription, RNA processing or RNA decay remained unclear (Pret and Searles 1991; Fridell and Searles 1994; Kuan et al. 2004; Kuan et al. 2009; Brewer-Jensen et al. 2016). SU(S)/ZC3H4 interacts with the conserved WDR82 protein (Brewer-Jensen et al. 2016; Austenaa et al. 2015), which supports recruitment of SU(S)/ZC3H4 to early elongation complexes through interactions with the Ser5-phosphorylated C-terminal domain of RNAPII (Bae et al. 2020; Lee and Skalnik 2008). WDR82 can also associate with the COMPASS/Set1 methyltransferase and the PP1-PNUTS phosphatase complex, but it remains unclear whether SET1 or PP1 activity contribute to Restrictor function (Austenaa et al. 2015; Hughes et al. 2023; Estell et al. 2023; Russo et al. 2023). SU(S)/ZC3H4 facilitates degradation of target RNAs (Kuan et al. 2009), with ZC3H4 recently shown to bind the adapter protein ARS2 for association with the Nuclear Exosome Targeting Complex (NEXT) (Rouvière et al. 2023; Estell et al. 2023). Restrictor has been postulated to promote RNAPII termination (Austenaa et al. 2021; Estell et al. 2023; Estell and West 2025). But, in contrast to both the CPA machinery and Integrator, Restrictor lacks any known catalytic activity, raising questions about the mechanisms of Restrictor function.

Restrictor’s target specificity also remains unclear, with several distinct models being proposed. Elegant genetic studies in Drosophila showed that sensitivity to SU(S) is conferred by the initially transcribed sequence and demonstrated that insertion of a consensus 5’ SS into a SU(S) target RNA prevents transcriptional suppression (Fridell and Searles 1994; Murray et al. 1997). In that work, the presence of a strong 5’ SS protected transcripts from targeting by Restrictor, explaining its preference for ncRNAs. However, work on mammalian Restrictor suggested several new and conflicting models for Restrictor targeting. Although some studies supported a conserved mechanism with Drosophila SU(S) (Estell et al. 2023), others hypothesized that Restrictor is selectively recruited to transcripts by the presence of weak 5’ SSs (Austenaa et al. 2021), or alternatively, that ZC3H4 activity is repressed at CpG islands, through competitive interactions of WDR82 with COMPASS/Set1 (Hughes et al. 2023). Distinguishing between these models through analysis of genomic data is complicated by the co-occurrence of many sequence and chromatin features at mRNAs. For example, genes originating from CpG islands often display high gene activity, contain strong 5’ SSs, and exhibit active histone modifications (Almada et al. 2013; Scruggs et al. 2015). A better understanding of how sequence drives target specificity of Restrictor requires experiments that can isolate the effect of sequence from confounding influences of chromatin and genomic context.

In this work, we use unbiased sequence screens to systematically probe Restrictor specificity in mouse embryonic stem cells (mESCs), demonstrating that Restrictor exhibits promiscuous activity that is selectively blocked by a strong 5’ SS, with little observed role for promoter sequence content. Further, using nascent RNA assays paired with rapid (< 1 hour) ZC3H4 degradation, we find that Restrictor acts broadly on uaRNAs but has very little activity on mRNAs, establishing Restrictor as a key determinant of transcription directionality at divergent promoters. We also find that the loss of Restrictor allows for the upregulation and extension of many uaRNAs and eRNAs, which can result in transcriptional interference and indirectly affect mRNA expression. Critically, we demonstrate that Restrictor acts directly on transcription by slowing RNAPII elongation over the first several kb. This Restrictor-mediated reduction in elongation rate makes RNAPII susceptible to early termination, including by the CPA machinery and Integrator. Despite lacking catalytic function, we report that Restrictor manipulates RNAPII activity to facilitate the termination of spurious RNAs, and safeguard mRNA expression.

## RESULTS

### Unbiased sequence screen highlights the importance of initially transcribed region in targeting of Restrictor

To comprehensively probe the determinants of Restrictor selectivity, we depleted Restrictor subunits ZC3H4 and WDR82 in mouse embryonic stem cells (mESCs) and screened transcription across thousands of integrated sequences using INSERT-seq. This approach measures the effects of a library of sequences that are integrated into the initially transcribed region of a ncRNA reporter locus (**Fig. 1A**). We recently demonstrated the power of this approach to define how the transcribed sequence impacts RNA output, comparing sequences derived from the 5’ ends of mouse mRNAs, lncRNAs, uaRNAs and eRNAs to a repertoire of synthetic sequences. Here, we leverage INSERT-seq to interrogate the effect of the transcribed sequence on Restrictor function, with all sequences tested at a specific genomic locus, with no variation in the promoter or chromatin context.

**Figure 1.**
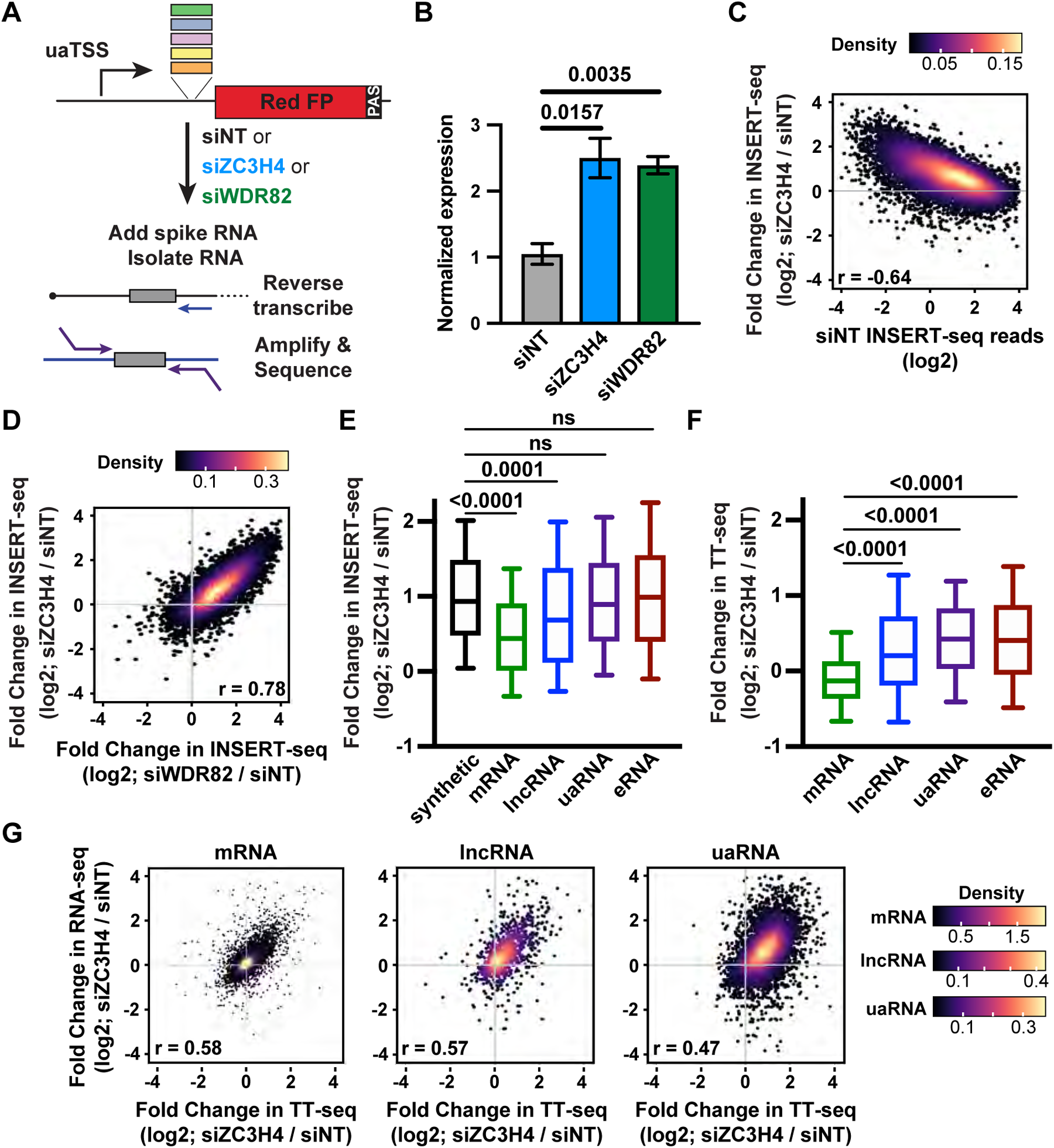
Sequences derived from mRNAs protect against Restrictor-mediated transcription repression. *(A)* Schematic of INSERT-seq. A library of 16,461 173-bp sequences was inserted at a uaRNA reporter locus in mESCs as described in Vlaming et al. (2022). This cell pool was treated with siRNAs against ZC3H4, WDR82 or a non-targeting control (NT) for 48 h. The effect of each inserted sequence on RNA expression was read out by high-throughput sequencing. *(B)* RT-qPCR measuring the level of RNA derived from the reporter locus in the library-containing cell pool. Total RNA was harvested 48 h after transfection with the indicated siRNAs. RNA levels were normalized to control gene TBP. Bars show mean and error bars report the standard deviation (siNT and siZC3H4, N = 3; siWDR82, N = 2). P-value from t-test. *(C)* Density scatter plot showing the fold change in INSERT-seq levels between siZC3H4 and siNT conditions, graphed against INSERT-seq expression levels for each sequence under siNT conditions (n = 9,334). *(D)* Density scatter plots comparing the fold change in INSERT-seq signal upon siWDR82 (x-axis) with siZC3H4 (y-axis) per inserted sequence (n = 9,334). *(E)* Box plots depicting the fold change in INSERT-seq signal after ZC3H4 KD for sequences derived from initially transcribed regions of various RNA classes (random synthetic sequences n = 906; mRNA n = 3609, lncRNA n = 308, uaRNA n = 1407, eRNA n = 1572). Boxplots have a line at the median with whiskers from 10-90th percentile. P-values from the Mann-Whitney test. ns = not significant, p > 0.05. *(F)* Box plots showing the fold change in TT-seq signal upon siZC3H4 at the endogenous genomic locations of the sequences shown in 1E. *(G)* Density scatter plots reporting the fold change in RNA-seq and TT-seq signal upon siZC3H4 for each RNA biotype (mRNAs n = 11960, lncRNAs n = 912 , uaRNAs n = 5798). For mRNAs and lncRNAs, RNA-seq and TT-seq reads were counted within exons. For uaRNAs, reads were counted from the uaTSS to 3 kb downstream. See also Supplemental Figure S1.

To test whether transcription at the reporter locus is normally attenuated by Restrictor, the cell pool containing our previously described sequence library inserted at the uaRNA reporter locus was transfected with siRNAs against ZC3H4, WDR82 or a non-targeting control (siNT) (**Supplemental Fig. S1A**), and RNA from was reverse transcribed for analysis by RT-qPCR. We observed a significant increase in reporter RNA abundance after both ZC3H4 and WDR82 knockdown (KD) as compared to siNT control, demonstrating that Restrictor broadly attenuates RNA synthesis at this locus (**Fig. 1B**). Samples in triplicate were spiked with in vitro transcribed RNAs to allow for absolute quantification of RNA levels and RNA produced from the reporter locus was amplified for sequencing (**Supplemental Fig. S1B**).

To evaluate the effect of Restrictor function on RNA levels across inserted sequences, we calculated the fold change in INSERT-seq signal between siZC3H4 or siWDR82 and siNT control conditions. In this comparison, sequences that are normally attenuated by Restrictor would have increased abundance following siZC3H4 or siWDR82 KD. Indeed, both ZC3H4 and WDR82 KD led to a broad increase in INSERT-seq signal (**Fig. 1C and Supplemental Fig. S1C**, respectively). The largest increases were observed for inserts with low expression in siNT cells, consistent with RNAs containing those sequences being repressed in the control condition, and in agreement with previous work showing that Restrictor targets typically exhibit low expression. Notably, our results imply that Restrictor acts on most but not all sequences, despite being controlled by the same promoter in the same chromatin environment, emphasizing that current models for Restrictor specificity warrant further exploration (Hughes et al. 2023; Austenaa et al. 2021).

Direct comparison of the fold changes in RNA abundance following WDR82 vs. ZC3H4 KD revealed the expected agreement between the two subunits of the Restrictor complex (**Fig. 1D**). Therefore, since WDR82 can participate in additional transcription regulatory complexes, we focused subsequent analyses on samples from ZC3H4 KD, which is thought to be uniquely present in Restrictor. We compared the effect of ZC3H4 KD on the abundance of RNA from synthetic sequences as compared to sequences derived from the initially transcribed regions of mRNAs, lncRNAs, uaRNAs or eRNAs. We found that sequences derived from all ncRNA classes and synthetic controls were more abundant in cells depleted of ZC3H4. However, regions derived from mRNAs were significantly less affected by loss of Restrictor (**Fig. 1E**), as reported for endogenous mRNA loci as compared to ncRNAs (Austenaa et al. 2015, 2021; Estell et al. 2021, 2023; Rouvière et al. 2023). Importantly, the effect of ZC3H4 KD on RNA levels is similar between random synthetic controls and sequences from uaRNAs or eRNAs, which argues against models wherein Restrictor is specifically targeted to ncRNAs to mediate transcription attenuation (Austenaa et al. 2021). Instead, our data supports a contrasting model wherein mRNA sequences themselves harbor elements that prevent Restrictor activity (Fridell and Searles 1994; Kuan et al. 2004; Brewer-Jensen et al. 2016; Estell and West 2025). Together, our results confirm that features of the initially transcribed sequence significantly impact Restrictor activity, even when isolated from their endogenous promoters and genomic locations.

### Restrictor attenuates RNA synthesis at non-coding loci

We performed transient transcriptome sequencing (TT-seq) to monitor newly synthesized RNA in mESCs after ZC3H4 KD (**Supplemental Fig. S1D**). To validate our findings from INSERT-seq, we first focused on the genomic regions that correspond to the initially transcribed sequences present in the INSERT-seq library (i.e. from the TSS +6 to +179). We observe that TT-seq signal from the early gene bodies of mRNAs is largely unchanged after ZC3H4 KD (**Fig. 1F**). In contrast, RNA synthesis within the same window downstream of TSSs for lncRNAs, uaRNAs and eRNAs is broadly increased after siZC3H4 (**Fig. 1F**).

We then defined differentially expressed RNAs in cells treated with siZC3H4, using TT-seq data within annotated genes. We observed a strong selectivity of Restrictor for ncRNAs, with very few mRNAs upregulated by siZC3H4 (263 of 12,009 total, or ∼2%), and a considerably larger fraction of lncRNAs upregulated (110 of 923 lncRNAs, or 12%, **Supplemental Fig. S1E**). Moreover, we observed a general upregulation of uaRNAs, with nearly 40% of uaRNAs considered significantly upregulated (2,128 of 5,867 total, or 36.3%, **Supplemental Fig. S1E**). Of note, fewer than 20 ncRNAs of any biotype investigated were downregulated following ZC3H4 loss. Importantly, for all RNA biotypes, we observed a good correlation between fold changes in TT-seq and RNA-seq signals after ZC3H4 KD (**Fig. 1G and Supplemental Fig. S1F**). The accumulation of uaRNAs and lncRNAs in the RNA-seq dataset indicates that the increased synthesis of these ncRNAs upon ZC3H4 KD results in elevated levels of stable RNA, supporting a general stabilization of non-coding RNA species in cells depleted of Restrictor (Kuan et al. 2009; Estell et al. 2023; Rouvière et al. 2023).

Altogether, our findings are consistent with previous work evaluating changes in RNA levels after Restrictor depletion and support several additional conclusions. First, the results of our comprehensive INSERT-seq screen, and the high degree of similarity of sequence behavior at the reporter locus as compared to endogenous gene loci, reveals that Restrictor specificity is primarily conferred by the initially transcribed RNA sequence, rather than features of the promoter or chromatin context. Second, we find that sequence elements within mRNAs specifically prevent Restrictor function, allowing protein coding genes to escape a general attenuation activity of Restrictor.

### 5’ Splice sites are protective against Restrictor-mediated attenuation

To identify sequence elements that could prevent Restrictor function, we searched for enriched motifs in mRNA sequences that were unchanged after ZC3H4 KD in INSERT-seq, as compared to mRNA inserts that were significantly upregulated after ZC3H4 KD. This analysis identified the 5’ SS as the only significantly enriched motif in Restrictor-resistant mRNA inserts (**Fig. 2A**). As noted above, prior reports have connected the 5’ SS motif with Restrictor activity, albeit with differing models (Fridell and Searles 1994; Austenaa et al. 2021; Estell et al. 2023). Moreover, whether the process of splicing is involved in attracting or repelling ZC3H4 activity remains unclear.

**Figure 2.**
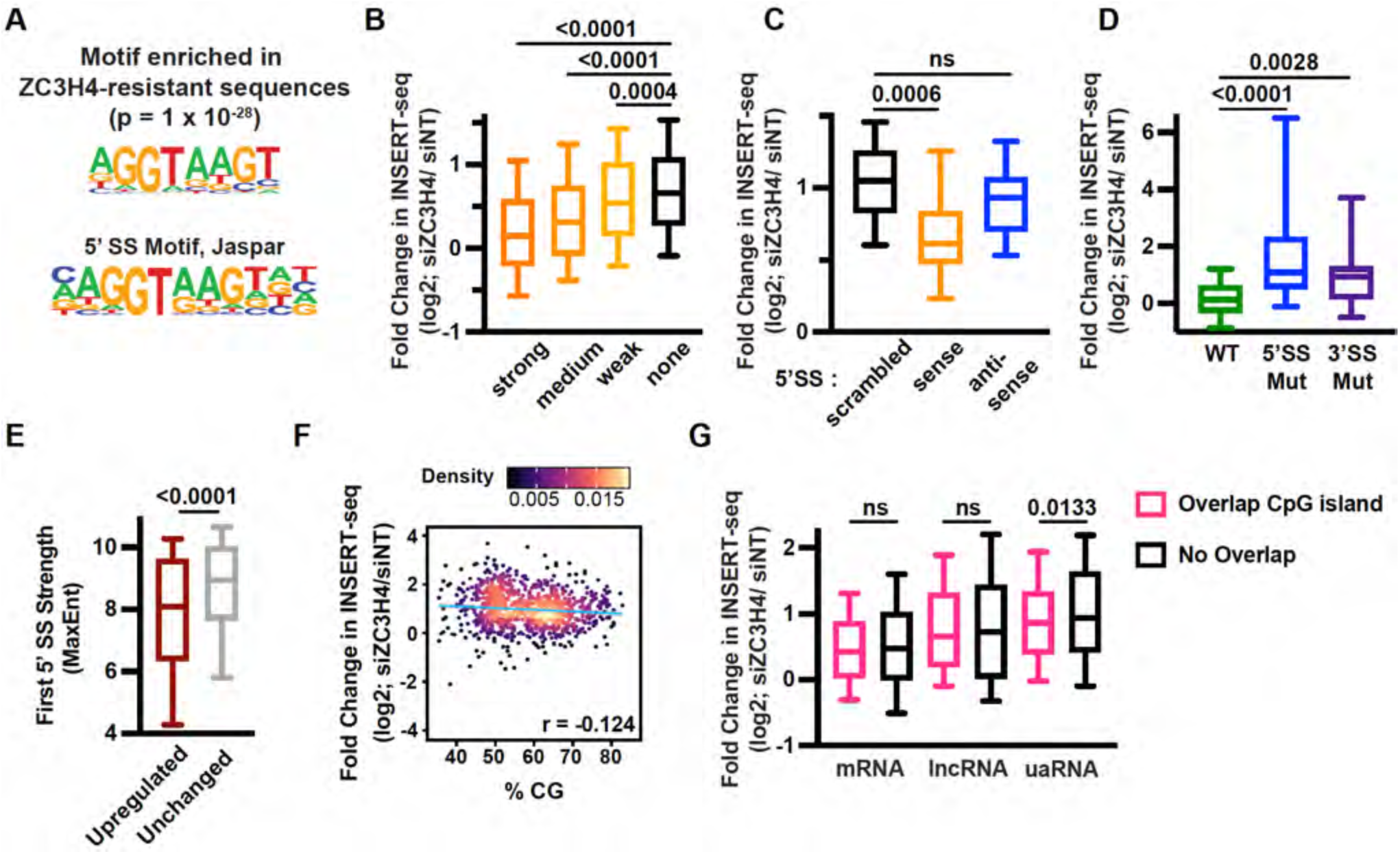
5’ splice sites, rather than CpG islands, confer resistance to Restrictor. *(A)* Motif enrichment in inserts derived from mRNAs unchanged by siZC3H4 in INSERT-seq (log2 fold change < |0.25|, n = 1,090) relative to sequences upregulated by siZC3H4 (log2 fold change > 0.5, n = 2,121), as determined by Homer. This motif has a similarity score of 0.96 to the 5’ SS motif from the Jaspar database (SD0001.1), which is shown for comparison. *(B)* Box plots reporting the fold change in INSERT-seq signal after ZC3H4 KD. Promoter-proximal inserts from mRNA sequences were separated based on maximum 5’ SS strength (MaxEnt) and compared to inserts that do not contain a 5’ SS (none, n = 1102, no match or MaxEnt <4). 5’ SSs were classified as in Almada et al., 2013, as strong (n = 1034, MaxEnt >8.77), medium (n = 595, MaxEnt 7.39-8.77) or weak (n = 878, MaxEnt 4-7.39). Box plots have a line at the median with whiskers from 10-90th percentile. Mann-Whitney test used to generate p-values. *(C)* Box plots as in B, for synthetic inserts where a scrambled (n = 18), sense (n = 38) or antisense (n = 18) 5’ SS was introduced into random background sequences. All 5’ SSs evaluated here were medium or strong (MaxEnt of > 7.39). Mann-Whitney test used to generate p-values. ns = not significant, p > 0.05. *(D)* Box plots showing fold change in INSERT-seq for sequences that contain either a splicing-competent intron, or the indicated mutant versions that are no longer spliced. (WT, n = 59; 5’ SS mut, n = 55; 3’ SS mut n = 28). Comparisons by Mann-Whitney test. *(E)* Box plots depicting the MaxEnt score (5’ SS strength) for first 5’ SSs at intron-containing mRNAs that are upregulated (n = 260) or unchanged mRNAs (n = 7,759) after siZC3H4. P-values from Mann-Whitney test. *(F)* Density scatter plot reporting the fold change in INSERT-seq after ZC3H4 KD with respect to % CG content. Data is shown for all synthetic inserts evaluated here (n = 906). *(G)* Box plot reporting the fold change in INSERT-seq signal after ZC3H4 KD for TSS-proximal inserts, separated based on overlap with a CpG island. Data is shown per biotype (left to right, n = 2918, 691, 129, 179, 819, 588). Mann-Whitney test used to generate p-values. ns = not significant, p > 0.05. See also Supplemental Figure S2.

Our INSERT-seq approach allowed us to disentangle the contribution of 5’ SS motif strength from other genomic features, by studying sequences with 5’ SS of varying strengths, all embedded within the same genomic context. We thus evaluated the change in INSERT-seq signal after ZC3H4 KD with respect to the strength of the 5’ SS motif. Investigation of sequences derived from the early gene bodies of mRNAs confirmed that inserts containing strong 5’ SS motifs (dark orange) were largely unaffected by ZC3H4 KD (**Fig. 2B**). As the strength of the 5’ SS motif decreased we observed stepwise increases in the effect of ZC3H4 KD on INSERT-seq signal (**Fig. 2B**). Importantly, mRNA sequences lacking a 5’ SS (black) were even more prominently affected by Restrictor than sequences with the weakest 5’ SS (yellow). These data indicate that a strong 5’ SS is protective from Restrictor and argue against weak a 5’ SS stimulating Restrictor recruitment or function.

To rule out context dependence, we assessed how the change in INSERT-seq signal upon siZC3H4 was influenced by 5’ SSs inserted into randomly generated background sequences. This was done in two ways; first we studied 5’ SS motif matches occurring in random sequences by chance, and second, we evaluated 5’ SSs specifically introduced into synthetic sequences (**Supplemental Fig. S2A and S2B,** respectively). In both instances, sequences containing strong 5’ SSs were less sensitive to loss of ZC3H4 than sequences with no or weak 5’ SSs. We then evaluated the effect of introducing 5’ SSs compared to introducing sequences with the same nucleotides in a scrambled order, or in the antisense orientation, both of which would be incompatible with U1 snRNP binding. Indeed, synthetic sequences with 5’ SS motifs inserted in the sense direction were significantly less affected by ZC3H4 KD than scrambled 5’ SSs (**Fig. 2C**). Furthermore, no significant differences were observed between inserts containing a scrambled vs. antisense 5’ SS (**Fig. 2C**), supporting that the 5’ SS sequence must be present on the RNA/non-template strand to block Restrictor activity.

Notably, most synthetic sequences with strong 5’ SSs inserted were not found to be spliced (3 out of 38 were spliced, and only at 10-20% splicing efficiency), consistent with the key determinant of Restrictor activity being the ability of the RNA to hybridize with U1 snRNP, rather than splicing competence (Fridell and Searles 1994). Indeed, when discarding inserts that showed any sign of splicing, inserted 5’ SSs still provide protection against Restrictor (**Fig. S2C**). Then, to determine whether the process of splicing also contributes to the protection from Restrictor activity, we mutated either the 5’ or 3’ SS within well-spliced introns and evaluated the changes in INSERT-seq signal after ZC3H4 KD. As anticipated, inserts containing a splicing-competent intron showed no significant changes in INSERT-seq signal after loss of ZC3H4 (**Fig. 2D**). In contrast, mutation of the 5’ SS within these introns led to a significant increase in INSERT-seq signal upon siZC3H4 (**Fig. 2D**), indicating that Restrictor was able to attenuate the expression of these sequences. Interestingly, inserts with 3’ SS mutations disrupting splicing were also increased in response to ZC3H4 knockdown (**Fig. 2D**), albeit with a smaller effect size. Surprisingly, these results indicate that disruption of splicing can also lessen the protection against Restrictor, even in the presence of an intact 5’ SS.

Our findings predict that the mRNA loci attenuated by Restrictor would have weaker 5’ SS consensus motifs than would mRNAs resistant to Restrictor activity. To test this model, we investigated the 5’ SS strength at mRNAs upregulated upon ZC3H4 depletion in our TT-seq experiment, as compared to mRNAs unchanged by Restrictor loss. Indeed, mRNAs upregulated by ZC3H4 depletion had weaker 5’ SSs than Restrictor-resistant, unchanged genes (**Fig. 2E**). We conclude that weak 5’ SSs do not recruit Restrictor, but instead that a strong 5’ SS can block promiscuous Restrictor-dependent transcription attenuation. Further, we find that the presence of a 5’ SS can inhibit Restrictor function independently of splicing, but that the process of splicing can contribute to protection against Restrictor activity.

### Neither CG nor CpG sequence content dictates Restrictor activity

Previous work has suggested a role for CpG islands in defining Restrictor target specificity and function, proposing that promoters embedded within CpG islands are less susceptible to gene attenuation by Restrictor. However, this model failed to explain the broad Restrictor sensitivity of uaRNAs which originate from the same CpG island as their partner mRNA TSS. Thus, to evaluate the influence of CG and CpG sequence content on Restrictor function, we compared the effects of ZC3H4 KD on INSERT-seq signals, with respect to sequence content of the investigated insert. Surprisingly, neither CG nor CpG content correlated well with changes in INSERT-seq reads following siZC3H4 (**Fig. 2F and Supplemental Fig. S2D**). To probe this phenomenon further, we separated sequences derived from mRNAs, lncRNAs and uaRNAs based on whether they overlap a CpG island (**Supplemental Fig. S2E**). We observed no significant differences in responsiveness to siZC3H4 between mRNA or lncRNA sequences that do vs. do not overlap a CpG island (**Fig. 2G**), with minimal differences in INSERT-seq signals observed for uaRNAs. Moreover, sequences derived from mRNAs respond differently to ZC3H4 KD than do ncRNAs, regardless of GC content. Thus, while the transcribed sequence itself is important, neither its CG nor CpG content serves as a major determinant of Restrictor’s ability to attenuate gene expression.

### Restrictor functions co-transcriptionally to attenuate gene expression

To confirm that we are measuring the direct targets of Restrictor, and to enable time-resolved assays following Restrictor depletion, we developed a system to rapidly degrade ZC3H4. The C-terminus of endogenous ZC3H4 was tagged with dTAG-2xHA-HiBiT, using CRISPR/Cas9 genome editing in mESCs (**Fig. 3A**), and three independent homozygous clones selected. Homozygous integration was confirmed by PCR and western blotting (**Fig. 3B and Supplemental Fig. S3A**). Based on the very rapid loss of the ZC3H4 protein upon treatment with dTAG-13 (hereafter referred to as dTAG, **Supplemental Fig. S3B**), all experiments were performed after a 40-minute final dTAG treatment. TT-seq libraries were generated after acute loss of ZC3H4, using a pulse of 4sU for the final 10 minutes, with spike-ins included to allow for absolute quantification. Replicate TT-seq libraries agreed well in both treatment conditions (**Supplemental Fig. S3C**), indicating agreement among the three homozygous ZC3H4-tagged clones.

**Figure 3.**
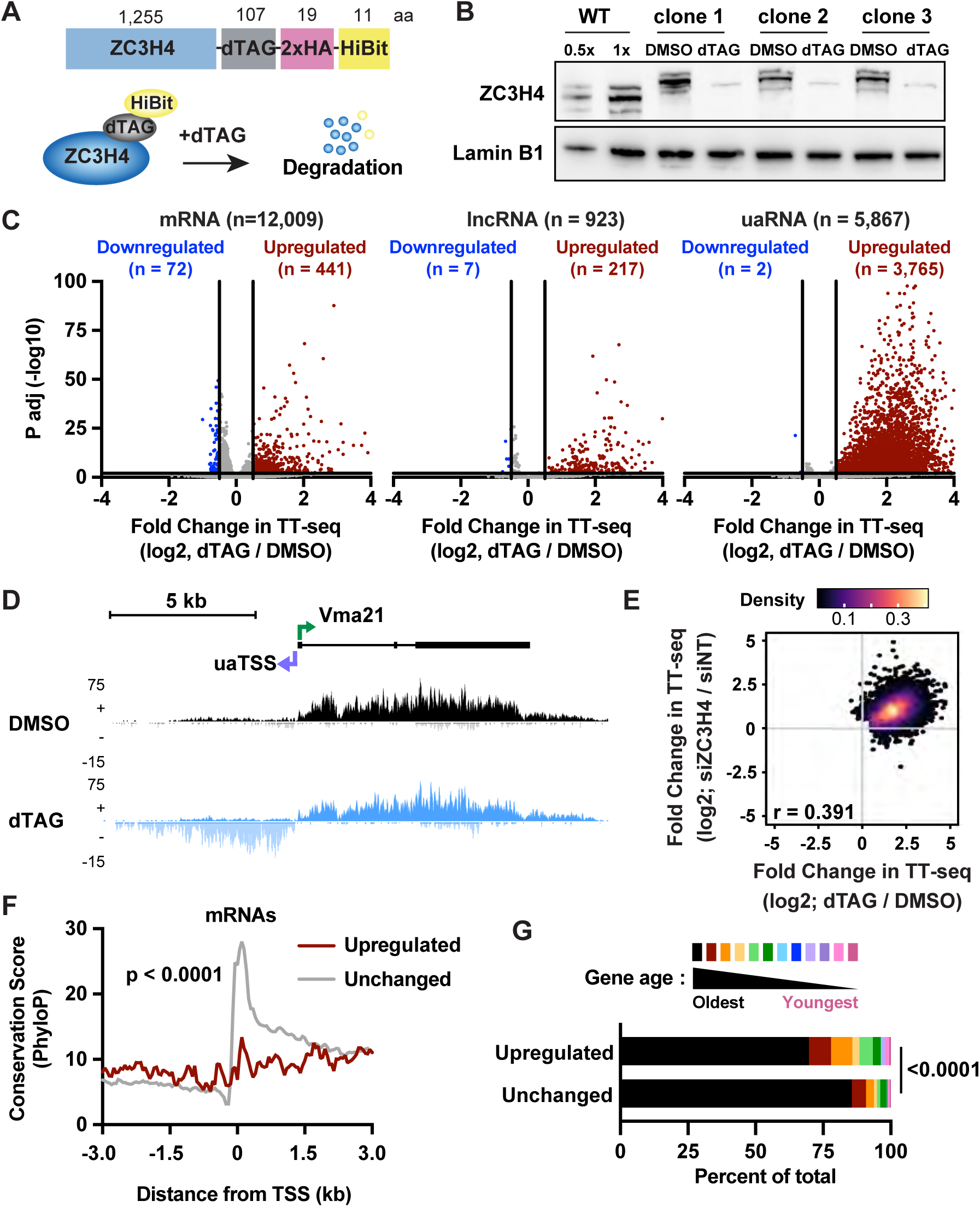
Restrictor selectively attenuates transcription of ncRNAs and poorly conserved mRNAs. *(A)* The C terminus of ZC3H4 was tagged with dTAG-2xHA-HiBiT. dTAG-13 (referred to as dTAG), induces degradation of the ZC3H4 fusion protein. *(B)* Western blots showing protein levels of ZC3H4 in the parental (WT) cell line and three independent clones with homozygously tagged ZC3H4. WT cells were untreated and were loaded at two concentrations. Tagged lines were treated with DMSO or dTAG for 40 minutes. *(C)* Volcano plots for each RNA biotype depict differentially expressed genes in dTAG vs. DMSO treated cells. For mRNAs and lncRNAs, TT-seq reads were calculated within exons. For uaRNAs, reads were counted from the uaTSS to 3 kb downstream. Affected genes were those with log2 fold change > 0.5 and p < 0.01. mRNAs with a log2 fold change < |0.25| were defined as unchanged (n = 7,547). *(D)* Stranded TT-seq signal in DMSO- and dTAG-treated conditions is shown at an example divergent promoter harboring an mRNA-uaRNA pair. *(E)* Density scatter plot comparing the fold change in TT-seq signal after dTAG-treatment (x-axis) with siZC3H4 (y-axis). Data is shown for mRNAs, lncRNAs or uaRNAs upregulated after either ZC3H4 depletion strategy (n = 4,773). *(F)* Metagene plot of PhyloP conservation scores at upregulated (n = 441) and unchanged (n=7,547) mRNAs, centered on the TSS and summed in 50-nt bins. P-value was generated using the Mann-Whitney test, comparing reads from TSS to +1 kb. *(G)* Bar plot depicts the distribution of gene ages (from Zhang et al, 2010) for upregulated and unchanged mRNAs. See also Supplemental Figure S3.

To quantify global effects of ZC3H4 loss on RNA synthesis, we defined differentially expressed genes between dTAG- and DMSO-treated conditions, generating separate volcano plots for mRNAs, lncRNAs and uaRNAs. As expected, most protein coding genes are unaffected by rapid loss of ZC3H4 (**Fig. 3C**; 3.7% of mRNAs are upregulated and 0.5% are downregulated upon dTAG treatment). In contrast, we observed a marked upregulation of ncRNA species. Specifically, 21.4% of lncRNAs, and 46.8% of uaRNAs are significantly upregulated after short-term depletion of ZC3H4 (**Fig. 3C**). The Vma21 mRNA-uaRNA divergent promoter is an example of this distinct behavior at coding vs. ncRNAs (**Fig. 3D**), with synthesis of the Vma21 mRNA unchanged after dTAG treatment, but with a significant increase in TT-seq signal at the associated uaRNA (**Fig. 3D**). These results are in line with the changes in gene expression observed after long-term loss of ZC3H4: graphing the fold change in TT-seq signal after ZC3H4 knockdown by siRNA vs. acute ZC3H4 degradation demonstrated good agreement between the two datasets (**Fig. 3E**). These data imply that many RNAs upregulated after long-term depletion of ZC3H4 are direct targets of ZC3H4 activity.

### Restrictor-resistant mRNAs exhibit higher sequence conservation

Our findings suggest that Restrictor-sensitive transcripts lack features that promote RNAPII elongation. To probe this idea, we investigated the sequence conservation across placental mammals around the set of mRNA promoters that were upregulated upon acute Restrictor depletion (**Fig. 3C**). Strikingly, the Restrictor-sensitive mRNA promoters exhibit low sequence conservation (**Fig. 3F**). By comparison, genes unaffected by Restrictor showed a strong peak of conservation extending from upstream of the TSS to ∼500 nt downstream (**Fig. 3F**). We wondered if the lower sequence conservation around Restrictor-sensitive gene 5’ ends was explained by shorter first exons at these genes, as introns have lower sequence conservation than exons. However, this was not the case, as the distance from TSS to first intron is moderately longer at upregulated mRNAs as compared to unchanged genes (**Supplemental Fig. S3D**). We thus examined the evolutionary ages of Restrictor-sensitive and Restrictor-resistant mRNAs. Notably, genes that were upregulated by Restrictor depletion were significantly younger in evolutionary age than unchanged genes (**Fig. 3G**). Thus, we propose that over evolutionary time, RNAs evolve features such as 5’ SS that render them less sensitive to attenuation by Restrictor.

### Loss of ZC3H4 allows for extended uaRNA and eRNA transcription that can interfere with expression of downstream mRNAs

Previous work has shown that transcription of uaRNAs and eRNAs is typically terminated within the first 1-2 kb, to prevent spurious RNA formation, interference with other transcribed units, and polymerase collisions (Cinghu et al. 2017; Flynn et al. 2016). Our data thus far broadly implicate Restrictor in suppressing synthesis of these ncRNAs, suggesting that loss of Restrictor might elicit transcriptional interference that impacts mRNA expression. To test this idea, we first generated heatmaps and composite metagenes of TT-seq signal from our control mESCs, aligned around all uaRNA TSSs significantly affected by Restrictor degradation (**Fig. 3C**, n=3,765). These data confirm that RNA synthesis does not generally extend more than 2 kb downstream of uaRNA TSSs (**Fig. 4A and 4B**). However, upon acute depletion of ZC3H4, both the amount of uaRNA synthesized and the length of the transcribed region were increased. Indeed, we find that uaRNA synthesis can persist well over 7.5 kb in the absence of Restrictor (**Fig. 4B and 4C**). Moreover, we noted that these extended uaRNAs could overlap nearby mRNA promoters, potentially influencing transcription initiation and/or elongation within these genes.

**Figure 4.**
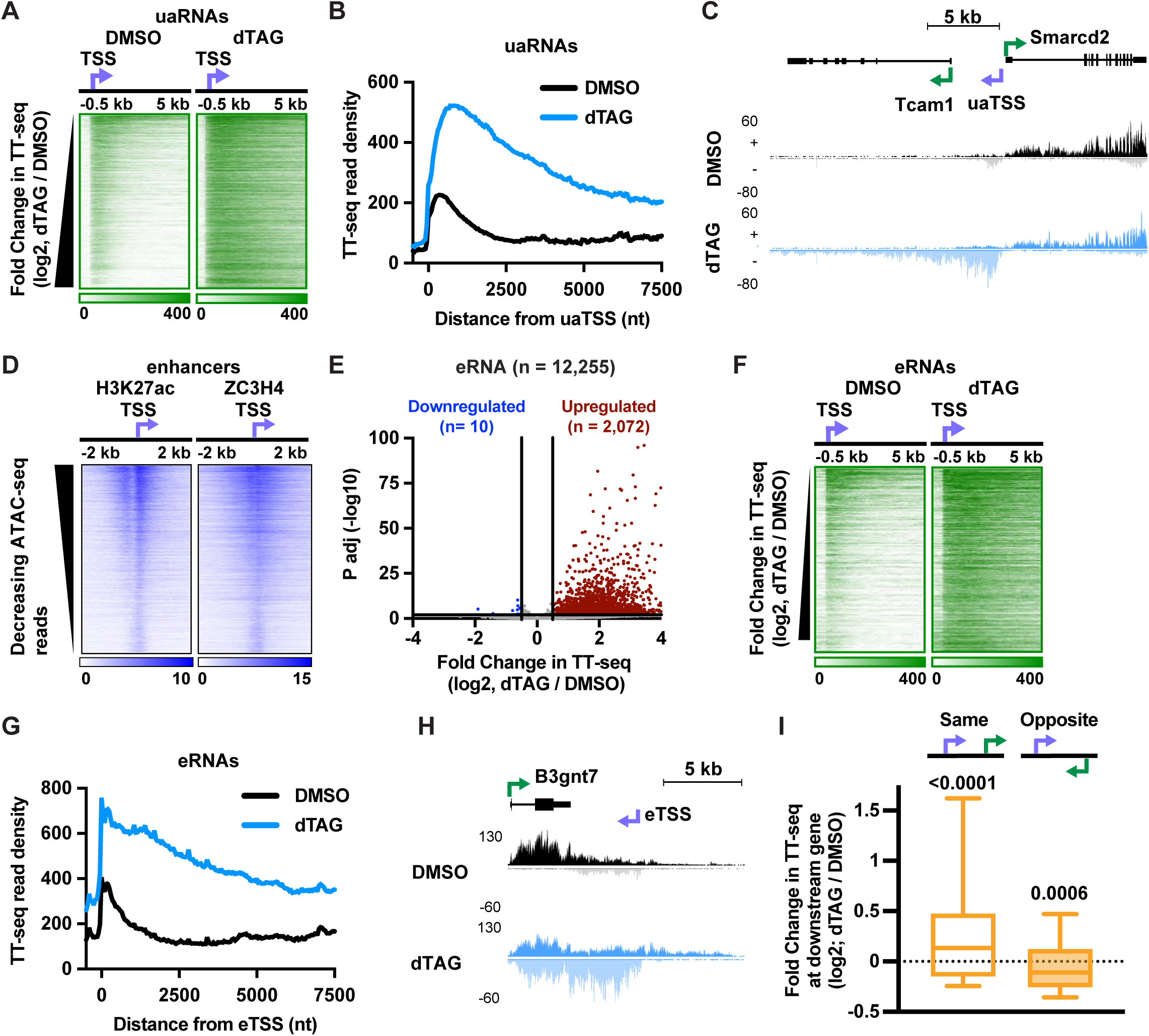
Loss of ZC3H4 allows for extended ncRNA transcription that can alter expression of downstream genes. *(A)* Heatmaps depicting the indicated TT-seq datasets at upregulated uaRNAs (n = 3,765), ranked by increasing fold change in TT-seq between DMSO and dTAG conditions. Data is aligned to the uaTSS and summed in 50-nt bins. *(B)* Metagene plots of TT-seq signal in DMSO- and dTAG-treated cells at upregulated uaRNAs (as in A). Data is aligned to the uaTSS, and summed in 50-nt bins. *(C)* Stranded TT-seq signal is shown at an example divergent promoter, where the extension of uaRNA synthesis after loss of ZC3H4 causes readthrough into the downstream mRNA gene, Tcam1. *(D)* Heatmaps depict ChIP-seq for H3K27 acetylation (from Vlaming et al. 2022) and ZC3H4 (from Hughes et al. 2023) at enhancers. Data is aligned to the dominant TSS within each enhancer (shown by arrow, n = 12,255). *(E)* Volcano plot depicting differentially expressed eRNAs in dTAG vs. DMSO-treated cells (log2 fold change > 0.5 and p < 0.01). Reads were summed from the eTSS to + 2 kb. *(F and G)* Same as A and B, but for upregulated eRNAs (n = 2,072). *(H)* Stranded TT-seq signal is shown for an example eRNA downstream and on the opposite strand of the B3gnt7 mRNA. The dominant TSS of the enhancer is indicated. *(I)* Box plots reporting the fold change in TT-seq upon ZC3H4 depletion for mRNAs within 5 kb downstream of upregulated uaRNAs or eRNAs. Data is separated based on the directionality of the uaRNA or eRNA with respect to the downstream mRNA (Same strand n = 145, Opposite strand n = 357). Box plots have a line at the median and whiskers depict 10-90^th^ percentiles. P-values calculated using the one sample Wilcoxon signed-rank test. See also Supplemental Figure S4.

We then investigated whether loss of Restrictor had similar effects on eRNA synthesis, given a report of Restrictor activity at super enhancers (Estell et al. 2021). We first identified putative enhancer loci using a combination of nascent RNA and chromatin accessibility data (**Supplemental Fig. 4**). The designation of these loci as putative enhancers was validated by the widespread presence of the active histone mark H3 Lysine 27 acetylation (H3K27ac) at these sites (**Fig. 4D**). Importantly, evaluation of previously generated ZC3H4 ChIP-seq in mESCs (Hughes et al. 2023) demonstrates a widespread occupancy of ZC3H4 at enhancers (**Fig. 4D**). Using TT-seq reads within 2 kb of the dominant TSS within each enhancer, we identified > 2000 enhancers with significantly increased eRNA synthesis in cells rapidly depleted of ZC3H4 (**Fig. 4E**). Heatmaps and composite metagenes of TT-seq signal at these loci confirmed both the short length and low-level expression of eRNAs in control cells, and the substantial increase in eRNA length and expression in cells depleted of ZC3H4 (**Fig. 4F and 4G**). As observed for uaRNAs, we found striking expansion of eRNAs in cells lacking Restrictor activity, with many extending > 7.5 kb (**Fig. 4G**). Further, we found many examples where continued synthesis of an eRNA overlapped an mRNA gene (e.g., **Fig. 4H**).

To investigate whether aberrant elongation of ncRNAs across mRNA genes had the potential to influence gene activity, we focused on mRNA genes that were within 5 kb downstream of uaRNAs or eRNAs upregulated in ZC3H4-depleted cells. Genes downstream of upregulated ncRNAs and on the same strand showed modestly increased TT-seq signal (across all annotated exons), suggesting that readthrough into these mRNA transcripts could increase their expression (**Fig. 4I, same**). Moreover, mRNAs on the opposite strand from uaRNAs or eRNAs upregulated upon ZC3H4 loss were significantly downregulated, demonstrating transcription interference (**Fig. 4I, opposite**). These data suggest that, despite not directly affecting the expression of many protein-coding genes, Restrictor activity indirectly safeguards mRNA expression, by preventing transcription interference. Further, we find that, in the absence of Restrictor activity in upstream antisense or enhancer regions, RNAPII is not efficiently terminated by other mechanisms or machineries (e.g., Integrator, CPA).

### Rapid depletion of ZC3H4 alters RNAPII elongation

To dissect how rapid depletion of ZC3H4 affects transcription elongation, we generated Precision Run-On (PRO)-seq libraries in DMSO- and dTAG-treated cells, using the same three clonal lines as above. PRO-seq involves the transcriptional incorporation of a single biotin-NTP, which is used to stringently isolate nascent RNAs and map the position of engaged RNAPII at single nucleotide resolution. Spike-ins were included to allow for absolute quantification across samples (**Supplemental Fig. S5A**). We first investigated the full set of mRNAs, lncRNAs and uaRNAs affected by ZC3H4 degradation in TT-seq data, comparing the fold changes between TT-seq and gene body PRO-seq signals. This analysis revealed good agreement between the two datasets (**Fig. 5A**), supporting that changes in gene expression after loss of ZC3H4 are due to altered RNAPII behavior.

**Figure 5.**
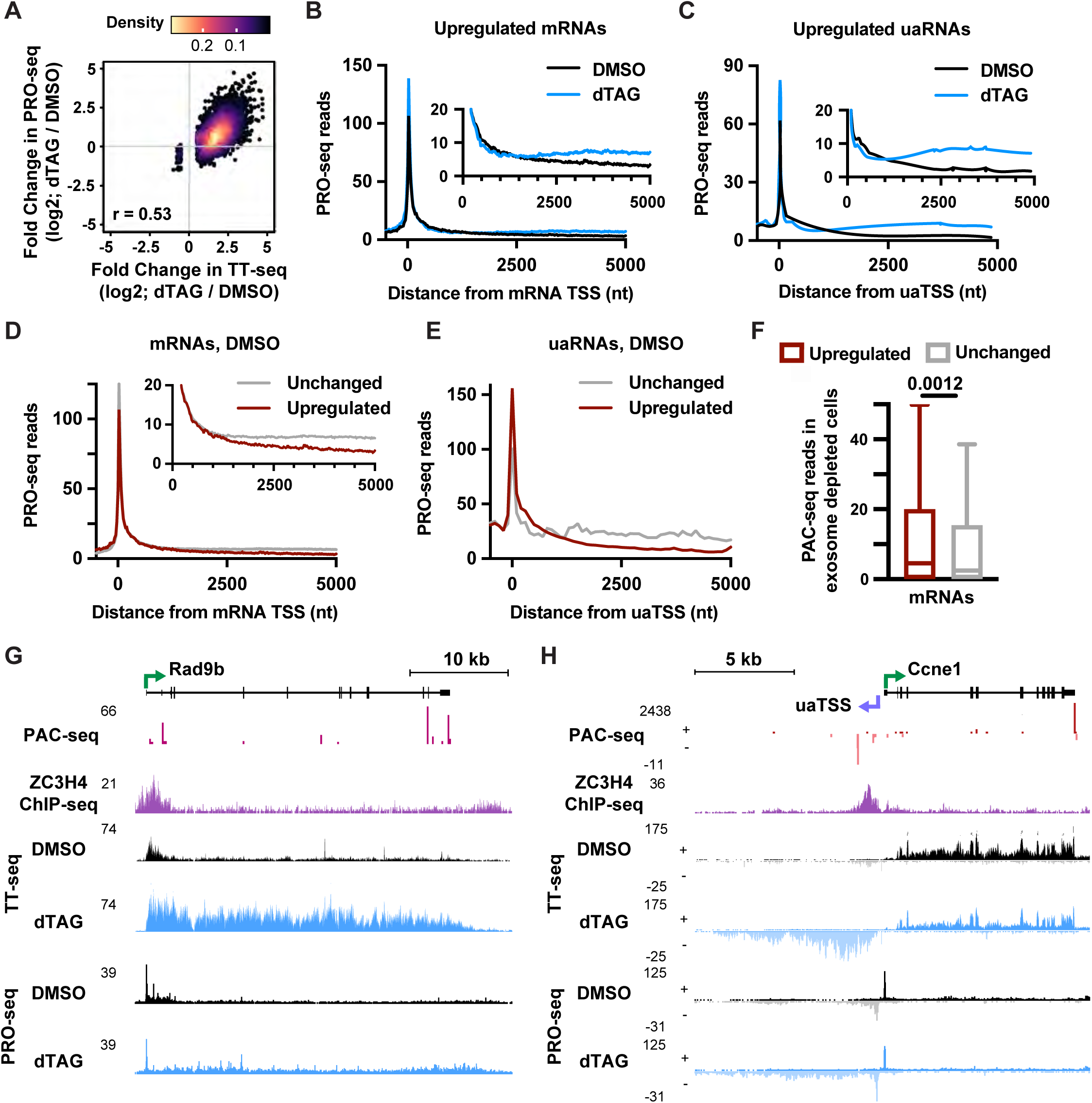
Rapid depletion of ZC3H4 enables productive elongation at genes where RNAPII is normally susceptible to premature termination. *(A)* Density scatter plot showing the fold change in reads upon ZC3H4 degradation, comparing data from TT-seq to gene body PRO-seq. Values are shown for differentially expressed mRNAs, lncRNAs and uaRNAs, as in Figure 3C. *(B)* Metagene plots of PRO-seq signal in DMSO- and dTAG-treated cells at upregulated mRNAs (n = 441). Data is aligned to the TSS, and summed in 25-nt bins. Inset highlights gene body PRO-seq signal. *(C)* Metagene plots of PRO-seq signal at upregulated uaRNAs, graphed as in (B), n = 3,765. *(D and E)* Metagene plots of PRO-seq signal at mRNAs (D) or uaRNAs (E), from DMSO-treated conditions. Data is aligned to the TSS, and summed in 25-nt bins for mRNAs and 100-nt bins for uaRNAs. Unchanged mRNAs, n = 7,547; uaRNAs, n=332. *(F)* Box plots report PAC-seq reads in cells depleted of the exosome (using siRRP40) to inhibit RNA decay. Reads were summed between TSS and + 3 kb at upregulated and unchanged mRNAs. Box plots have a line at the median and whiskers depicting 1.5 times the interquartile range. P-value was generated using the Mann-Whitney test. *(G)* Example mRNA gene Rad9b is upregulated after loss of ZC3H4. Data on the sense strand is shown for indicated data types and conditions. *(H)* Indicated datasets are shown at the Ccne1 divergent promoter. The Ccne1 mRNA is unchanged and the corresponding uaRNA is upregulated after loss of ZC3H4. Data is shown on both the sense (+) and antisense (-) strand. See also Supplemental Figure S5.

We next sought to define how transcription is changed upon rapid depletion of ZC3H4. We generated metagene plots of PRO-seq signal at genes upregulated upon acute loss of ZC3H4, comparing RNAPII profiles between DMSO and dTAG conditions. This analysis revealed a prominent peak of paused RNAPII at upregulated mRNAs, uaRNAs and lncRNAs (**Fig. 5B, 5C and Supplemental Fig. S5B**, respectively), that was increased in the absence of ZC3H4. Notably, promoter-proximal PRO-seq signal was significantly increased at all genes, including those unchanged by loss of ZC3H4 (**Supplemental Fig. S5C**), suggesting a broad elevation in paused RNAPII upon loss of Restrictor that is not connected to changes in RNA expression. Instead, the selective and significant increase in PRO-seq signal at genes upregulated upon ZC3H4 loss started ∼1 kb downstream of the TSS (**Fig. 5B, 5C, Supplemental Fig. S5B and S5D)**. These data indicate that ZC3H4 has a negative impact on productively elongating RNAPII at a subset of mRNAs, as well as many uaRNAs and lncRNAs.

To understand the mechanism of ZC3H4-mediated transcription repression, we investigated RNAPII distribution at target genes in control cells (**Fig. 5D and 5E**, DMSO conditions). Comparing ZC3H4-sensitive genes to those unchanged by degradation of ZC3H4, we observed that both sets of promoters exhibit similar, prominent pausing of RNAPII. However, after pause release and entry into productive elongation, PRO-seq signal at unchanged mRNAs and uaRNAs reached a plateau around 1 kb downstream, suggesting that RNAPII reaches stable, productive elongation at this point, which persists downstream. In contrast, the PRO-seq signal at ZC3H4-sensitive loci steadily declined across the first 5 kb of the gene body, never appearing to stabilize (**Fig. 5D and 5E**). These results imply continued attrition of elongating RNAPII across ZC3H4-sensitive genes in the presence of ZC3H4, consistent with premature termination. Critically, ZC3H4 depletion prevented the attrition of PRO-seq signal within this gene window (i.e. downstream of ∼1 kb, **Fig. 5B and 5C**), allowing RNAPII levels to flatten out or even increase within gene bodies. These data suggest that ZC3H4 represses transcription of target mRNAs and uaRNAs, likely through destabilization or decreased processivity of the RNAPII elongation complex.

### Transcription attrition at ZC3H4 targets generates short, polyadenylated RNAs

Although Restrictor has been suggested to drive transcription termination (Estell et al. 2023), it lacks any known catalytic activity. Therefore, it remains unclear how Restrictor might suppress transcription, and whether Restrictor might work together with the CPA machinery at cryptic PASs. To probe whether Restrictor might promote RNA cleavage and polyadenylation, as has been suggested for Drosophila SU(S) (Brewer-Jensen et al. 2016), we identified RNAs with polyA tails using Poly-A Click (PAC)-seq (Mimoso and Adelman 2023). Because such RNAs could be susceptible to degradation by the RNA exosome, PAC-seq was performed in cells treated with siRNAs against the exosome subunit RRP40, to stabilize these termination products. Evaluation of PAC-seq signal in siRRP40-treated cells revealed significantly more reads at ZC3H4-sensitive mRNAs as compared to those unchanged by ZC3H4 depletion (**Fig. 5F**). Elevated PAC-seq signal was detected at upregulated uaRNAs and lncRNAs as well (**Supplemental Fig. S5E**). These data reveal some of the termination products at Restrictor-sensitive loci are polyadenylated, implicating the CPA complex. Example Restrictor-sensitive mRNAs (**Fig. 5G and Supplemental Fig. S5F)** and uaRNAs (**Fig. 5H)** show distinct sites of PAC-seq reads in early gene bodies, near peaks of Restrictor ChIP-seq signal. These sites are coincident with the locations of drop-off in both TT-seq and PRO-seq signal in control cells and depletion of Restrictor allows RNAPII to proceed beyond these sites. Our data thus support that termination of some Restrictor-sensitive RNAs is mediated by the CPA machinery.

### Nearly half of all Restrictor-sensitive mRNAs are also targets of Integrator

To investigate a potential relationship between Restrictor and Integrator, we directly compared their gene targets in mESCs, using the changes in PRO-seq signal following acute degradation of ZC3H4 (this study) or INTS11 (Stein et al. 2022). We noted that far more mRNAs are affected by loss of INTS11 than ZC3H4, consistent with the notion that Restrictor primarily targets ncRNAs (**Fig. 6A**, INTS11-affected n= 2,093; ZC3H4-affected n= 835). We quantified the changes in PRO-seq in the gene body of all affected mRNAs and clustered genes into four groups using k-means clustering. This analysis revealed three main groups: genes upregulated after loss of either ZC3H4 or INTS11 (Cluster 1, n = 378), genes that are upregulated only after loss of ZC3H4 (Cluster 2, n = 457) or only after loss of INTS11 (Clusters 3 and 4, n = 1,715) (**Fig. 6A**). We note that 45% of ZC3H4-sensitive genes are also INTS11 targets, with the overlap between the two groups much higher than expected by chance (Fisher test, odds ratio 4.3, p < 2e-16). Notably, this level of overlap was not apparent when comparing RNA following long-term knockdown of ZC3H4 and INTS1 (Estell et al. 2021), underscoring the utility of acute protein degradation approaches. This finding suggests that transcription suppression by Restrictor involves Integrator-mediated termination at some loci. A majority of Integrator targets, however, did not show a sensitivity to Restrictor loss, supporting that Integrator can terminate transcription independently (Wagner et al. 2023). Indeed, browser shots of individual genes that are consistent targets of Integrator, such as Jun (**Fig. 6B**, Gardini et al. 2014; Stein et al. 2022), confirmed that loss Restrictor did not affect their transcription.

**Figure 6.**
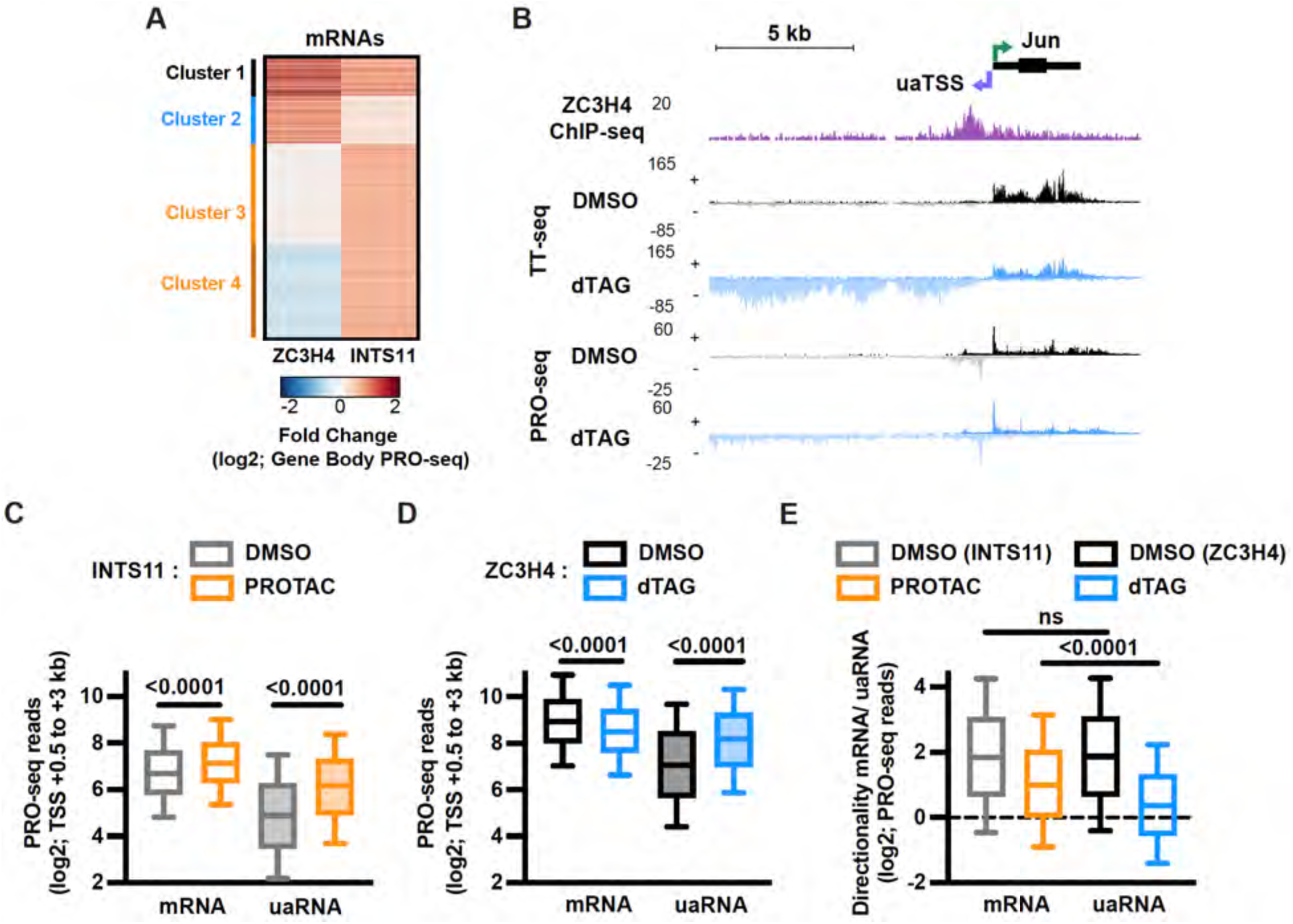
Restrictor is the primary determinant of transcription directionality at promoters. *(A)* Heatmap depicting the fold change in gene body PRO-seq signal after rapid depletion of ZC3H4 (dTAG, this study) or INTS11 (PROTAC, Data from Stein et al., 2022) as compared to control (DMSO). Data is shown for mRNAs with at least 5 gene body PRO-seq reads in all conditions (11,476 genes total) and a log2 fold change > 0.5 in gene body PRO-seq signal after loss of ZC3H4 or INTS11 (n = 2,550). Genes were classified into 4 clusters by k-means clustering Cluster 1, n = 378; Cluster 2, n = 457; Clusters 3 and 4, n = 1,715). *(B)* Indicated datasets are shown at the divergent Jun promoter. The Jun mRNA is unchanged but the corresponding uaRNA is upregulated after loss of ZC3H4. *(C)* Box plot reporting PRO-seq reads downstream of mRNA and uaRNA TSSs in control (DMSO) and INTS11-depleted (PROTAC) cells. PRO-seq reads were counted from the TSS +0.5 to + 3 kb. Box plots have a line at the median and whiskers depict 10-90th percentiles. P-values were generated using the Wilcoxon test. *(D)* Same as C, except for control (DMSO) and ZC3H4-depleted (dTAG) cells. *(E)* The Directionality score was calculated by dividing gene body PRO-seq reads at mRNAs over uaRNAs. Box plot reports the distribution of directionality scores per indicated condition. P-values were generated using the Wilcoxon test.

### Restrictor is a key mediator of transcriptional directionality at divergent promoters

Our investigation of Restrictor versus Integrator activity at mRNA promoters emphasized the preference of Restrictor for targeting uaRNAs. For example, we noted that ZC3H4 ChIP-seq signal was present near the uaRNA partner of Jun (**Fig. 6B**), and that antisense transcription was substantially increased upon Restrictor loss. We thus compared the effects of Restrictor and Integrator on the directionality of transcription at divergent promoters. In line with previous work, we find that depletion of INTS11 causes a broad increase in early elongation complexes across both protein coding genes and the associated upstream antisense RNAs (Lykke-Andersen et al. 2021; Stein et al. 2022). Quantification of PRO-seq signal in the window from 500 to 3000 nucleotides downstream of the TSSs shows significant increases upon INTS11 degradation, at both mRNAs and uaRNAs (**Fig. 6C**). In contrast, ZC3H4 causes a widespread increase in PRO-seq reads in the uaRNA direction, but no increase in the sense direction – with even a decrease in PRO-seq reads observed upon ZC3H4 loss (**Fig. 6D**). Thus, when we calculate a Directionality Index that compares PRO-seq signal within 3 kb of divergent mRNA-uaRNA promoters, we find that both Integrator and ZC3H4 depletion substantially reduce directionality (**Fig. 6E**). However, given the selectivity of Restrictor towards repression of uaRNAs, loss of Restrictor causes a significantly larger change in directionality, with the sense and anti-sense reads (mRNA and uaRNA) becoming nearly equivalent in cells lacking Restrictor (**Fig. 6E**). We conclude that Restrictor is a central determinant of transcription directionality.

### ZC3H4 suppresses uaRNA transcription by reducing RNAPII elongation rate

The broad upregulation of uaRNA synthesis in cells rapidly depleted of ZC3H4 or INTS11, and our acquisition of both PRO-seq and TT-seq data in mESCs under these conditions, allowed us to compare the mechanisms of action of Restrictor versus Integrator. First, we investigated the PRO-seq signal at all uaRNAs in both control and factor-depleted datasets. As shown for the set of upregulated uaRNAs in **Fig. 3C**, ZC3H4 degradation led to a general increase in PRO-seq signal at uaRNAs, starting around +1 kb, and persisting for many kb downstream (**Fig. 7A**). In contrast, INTS11 degradation caused a more marked increase in promoter-proximal PRO-seq signal, consistent with Integrator selectively targeting paused RNAPII. Loss of Integrator-mediated termination allows paused RNAPII to escape into gene bodies (**Fig. 7B**). However, consistent with previous work, we find that the aberrantly escaped polymerase is not productive and is susceptible to termination by other machineries (Lykke-Andersen et al. 2021; Stein et al. 2022; Hu et al. 2023; Blears et al. 2024; Cacioppo et al. 2024), such that the PRO-seq signal returns to baseline within several kb (**Fig. 7B**). Analysis of TT-seq signal at these loci confirmed that the stimulation of uaRNA synthesis upon INTS11 degradation only persists for several kb, whereas ZC3H4 degradation allowed for much longer extension of these RNAs (**Supplemental Fig. S6A**).

**Figure 7.**
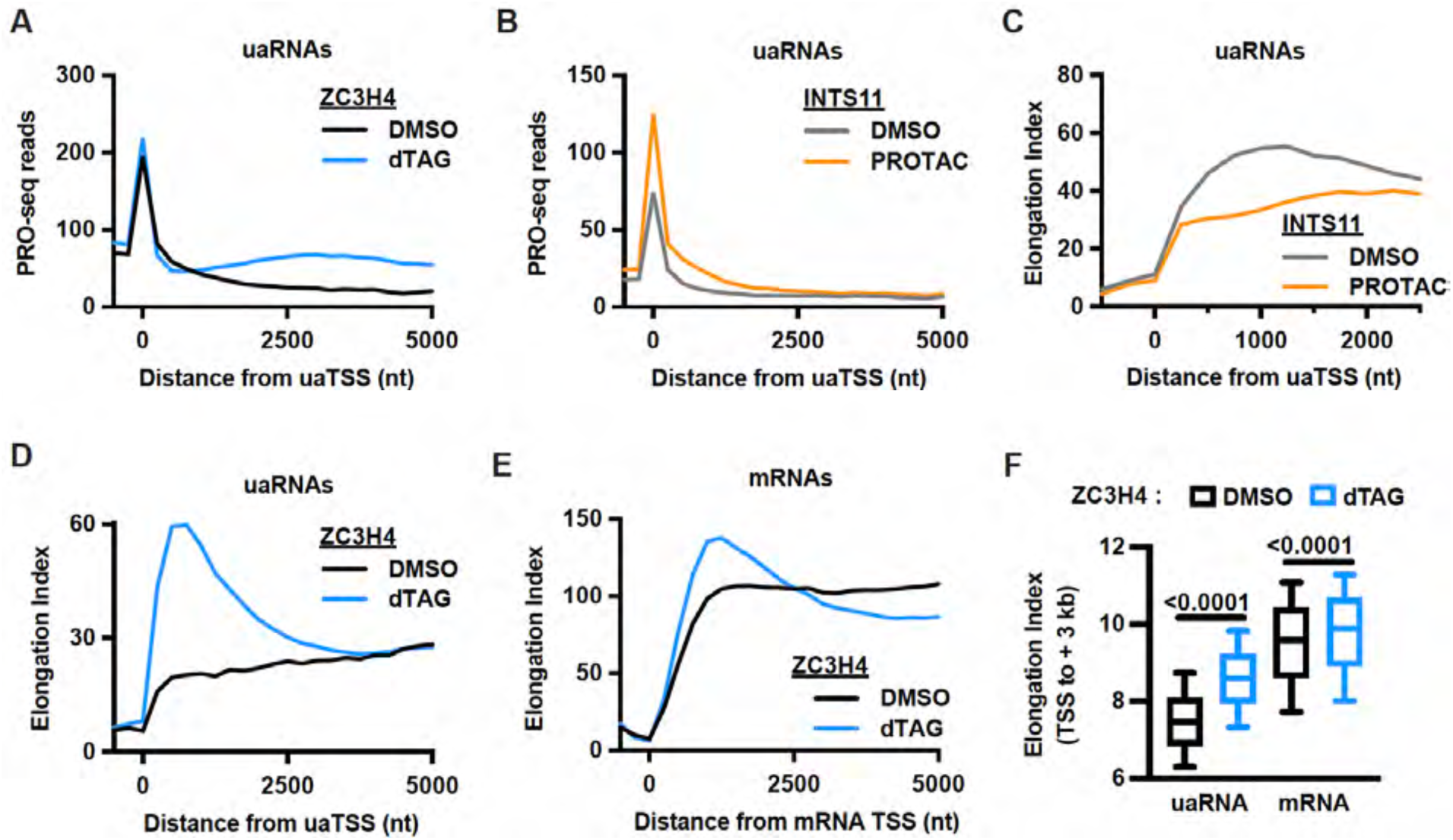
Restrictor reduces RNAPII elongation rate. *(A)* Metagene plots of PRO-seq signal in ZC3H4-tagged cells after dTAG or DMSO treatment at uaRNAs (n = 5,867). Data is aligned to the uaTSS, and summed in 250-nt bins. *(B)* Same as A, except for INTS11-tagged cells after PROTAC or DMSO treatment. *(C)* Metagene plots of elongation index at uaRNAs in PROTAC- and DMSO-treated INTS11-tagged cells. Elongation index was calculated as the ratio of TT-seq signal (RNA synthesis) to PRO-seq signal (elongating RNAPII) per 250-nt bin. x-axis was truncated at + 2.5 kb due to PRO-seq signal falling below threshold at this point in both datasets. *(D)* Elongation index metagenes at uaRNAs for ZC3H4-tagged cells after dTAG- or DMSO-treatment. *(E)* Elongation index metagenes aligned to mRNA TSS (n = 12,009) per 250-nt bin. *(F)* Box plot reporting the sum of elongation index values from TSS to +3 kb in dTAG- and DMSO-treated ZC3H4 cells at uaRNAs (n = 5,867) and mRNAs (n = 12,009). Box plots have a line at median with whiskers from 10-90th percentile. See also Supplemental Figure S6.

Mechanistically, we and others have shown that RNAPII that escape mRNA promoters precociously in Integrator-deficient cells exhibit elongation defects, with a failure to attain the normal elongation rates observed in control cells (Stein et al. 2022; Hu et al. 2023). To evaluate this at uaRNAs, we inferred elongation rate by dividing the signal for RNA synthesis (TT-seq) by the signal representing RNAPII density (PRO-seq), as performed previously (Žumer et al. 2021; Stein et al. 2022; Mimoso and Adelman 2023). These data confirmed that after pause release, RNAPII accelerates as it transcribes across uaRNAs in control cells (**Fig. 7C**, gray). However, this increase in elongation index is dampened in cells lacking INTS11 (**Fig. 7C**, orange). Strikingly, performing the same analysis on control and ZC3H4-depleted cells reveals a very different effect, with elongation index strongly increasing upon Restrictor loss (**Fig. 7D**). This apparent increase in elongation rate in cell lacking ZC3H4 suggests that Restrictor normally slows elongation across uaRNAs.

### Restrictor broadly slows early transcription elongation

To more fully evaluate the model that Restrictor reduces elongation rate, we calculated the elongation index in control and ZC3H4-depleted cells at mRNAs and lncRNAs (**Fig. 7E and Supplemental Fig. S6B**). At both RNA biotypes, we observe that RNAPII increases its apparent elongation rate over the first several kb in control cells, as anticipated (Jonkers et al. 2014; Fong et al. 2022). This acceleration is increased in cells lacking ZC3H4, suggesting that Restrictor can modulate RNAPII elongation rate at mRNA and lncRNA loci as well. Indeed, the increase of elongation index within the first several kb upon ZC3H4 loss is significant across all RNA biotypes investigated (**Fig. 7F and Supplemental Fig. S6C**). Together, our data support that ZC3H4 functions by reducing elongation rate, thereby facilitating termination by other mechanisms and machineries.

## DISCUSSION

Our results build on classical studies of SU(S) in *Drosophila* and more recent investigations of mammalian ZC3H4 to provide a revised model for ZC3H4 activity. We find that Restrictor is central to RNA quality control, both preventing transcriptional interference and supporting the directionality of transcription at mammalian promoters. In addition to a widespread role for Restrictor in suppressing upstream antisense transcription, we uncover a broad effect of Restrictor on the synthesis of eRNAs. Based on our findings, we propose the following model for Restrictor activity: Restrictor associates globally with RNAPII in early elongation, as the nascent RNA is sufficiently extended to allow for binding by ZC3H4. ZC3H4 would associate with RNA through some combination of its arginine-rich and zinc-finger motifs (Murray et al. 1997; Turnage et al. 2000; Estell et al. 2023), with interactions stabilized by WDR82 association with Ser5-P RNAPII (Bae et al. 2020; Lee and Skalnik 2008). Restrictor reduces transcription elongation rate, making RNAPII susceptible to termination and causing transcription attenuation within several kb from the TSS. However, RNAs that contain 5’ SSs in their initially transcribed regions, including the vast majority of mRNAs, will be bound by the U1 snRNP to relieve the suppression by Restrictor.

This model explains the broad activity of Restrictor on ncRNAs and seems parsimonious from an evolutionary perspective: new or spurious transcription start sites can arise anywhere in the genome and Restrictor’s default behavior would be to suppress the expression of RNAs from these sites. Indeed, our work reveals that ZC3H4 widely occupies enhancers and significantly reduces eRNA synthesis at ∼20% of these loci. We envision that, as transcription units evolve, they will gain sequences such as 5’ SSs, to escape Restrictor activity. We present evidence that Restrictor can work with the CPA machinery and Integrator, however slower elongation could sensitize transcripts to other modes of termination, including RNAPII degradation after irreversible stalling (Aoi et al. 2021; Reese 2023; Noe Gonzalez et al. 2021) or DNA-dependent termination (e.g., T-tracts, Han et al. 2023; Davidson et al. 2024). Notably, our previous study indicated that RNAPII is termination-prone in the absence of positive signals (Vlaming et al. 2022), and the promiscuous activity of Restrictor at ncRNA loci offers a potential explanation for this behavior.

Our work demonstrates that the initially transcribed sequence is the critical determinant of target specificity. Using a screen of thousands of sequences derived from various coding and non-coding RNAs, as well as synthetic, designed sequences, we demonstrate that a consensus 5’ SS prevents Restrictor activity, independently of splicing. Importantly, sequences with 5’ SS are specifically insensitive to Restrictor, rather than sequences lacking 5’ SS being particularly sensitive. We interpret this as supporting a general interaction of Restrictor with early elongation complexes that is disrupted by the recruitment of the U1 snRNP.

Transcription directionality has been largely attributed to the U1-PAS axis, since the initially transcribed regions of mRNAs are enriched for 5’ SSs, while uaRNAs contain more PAS motifs (Venters et al. 2019). However, evidence does not support the CPA machinery as the dominant termination complex at uaRNAs (Lykke-Andersen et al. 2021), implying that other termination machineries play bigger roles in suppressing uaRNA synthesis. Here, we find that Restrictor is a key contributor to transcription directionality, whose activity is prevented by 5’ SS. We thus propose that the driver of directionality is actually a U1-Restrictor axis, where Restrictor-mediated suppression of ncRNA transcription includes, but is not limited to, cleavage and polyadenylation at cryptic PASs.

In conclusion, we propose that Restrictor promiscuously binds RNAPII in early elongation to promote termination, with most mRNAs selectively protected from this activity by the presence of 5’ SS. We suggest that Restrictor activity is critical in mammalian cells (Su et al. 2021) due to its central role in preventing spurious transcription, driving the directionality of transcription at promoters and averting transcription interference. Our results place Restrictor in a new light: revealing that it sensitizes RNAPII for termination by slowing elongation, rather than acting as a termination machinery. It will be exciting in the future to biochemically and structurally dissect the mechanisms of Restrictor activity.

### Limitations

Our results do not support a role for promoter sequence or CG content in defining Restrictor activity. Although our findings differ from a recent model that suggested CpG island promoter conferred resistance to Restrictor (Hughes et al. 2023), our data are in line with early work from Drosophila, in which substitution of promoter sequences did not impact the effects of Su(S)/ ZC3H4 (Fridell and Searles 1994), but replacing a weak 5’ SS with a consensus 5’ SS eliminated gene responsiveness to Su(S) activity. Nonetheless, it remains possible that CG content or CpG islands contribute to Restrictor specificity in a manner that wasn’t measured in our assays.

## METHODS

Methods can be found at the end of this file.

## ACKNOWLEDGEMENTS

We thank all members of the Adelman lab, and Lillie Searles, whose elegant work on SU(S) provided the foundation for this work. This work was supported by the National Institutes of Health (NIH R01GM139960 to K.A.). C.A.M was supported by the Sophia H.Y Chang Fellowship, and H.V. was supported by the Human Frontier Science Program (LT000651/2018-L).

## AUTHOR CONTRIBUTIONS

Investigation: CAM, HV, NPdW KA. Conceptualization: CAM, HV, KA. Methodology: CAM, HV, KA Formal data analysis: CAM, HV Visualization: CAM, HV, KA Project administration: KA Supervision KA. Funding acquisition: KA Writing – original draft CAM, HV, KA Writing – review & editing: CAM, HV, KA

## MATERIALS AND METHODS

### Cell Culture

mESCs (CAST/129 hybrid background, female) were cultured on gelatin. For most experiments, KnockOut DMEM (ThermoFisher, 10829018) was used, supplemented with 15% KO serum replacement (ThermoFisher, 10828028), 1X penicillin-streptomycin (MP, TMS-005-C), 1X non-essential amino acids (MP, TMS-001-C), 1% β-ME (MP, ES-007-E), 1X GlutaMAX (ThermoFisher, 35050061), 1000 U/ml LIF (Cell Guidance Systems, GFM200), 1 µM MEK inhibitor (Stemgent, PD0325901), and 3 µM GSK3 inhibitor (Stemgent, CHIR99021). For the TT-seq and RNA-seq experiments after knockdown, wild-type F121-9 cells were grown in serum-free embryonic stem cell (SFES) medium with the same concentrations of LIF and inhibitors. SFES was composed of 50/50 NeuroBasal medium (Gibco) and DMEM/F12 medium (Gibco), supplemented with 0.5x B-27 (Gibco), 0.5× N-2 (Gibco), 2 mM l-glutamine (Gibco), 0.05% bovine albumin fraction V (Gibco) and 1.5 × 10−4 M monothioglycerol (Sigma). mESCs were cultured at 37°C with 5% CO_2,_ fed daily, passaged every two days.

S2 (*Drosophila*) cells were used to generate spike-ins for TT-seq and PRO-seq (as described below). S2 cells were cultured in Shields and Sang M3 medium (Sigma, S3652) supplemented with yeast extract (Sigma, Y-1000), bactopeptone (Difco, 211677) and 10% FBS (Thermo, 1600044). S2 cells were cultured at 27°C.

mESCs and S2 cells were routinely tested for mycoplasma contamination.

### RNAi

Cells were transfected with 30 nM siRNAs, using the RNAiMAX reagent (Invitrogen) in a suspension transfection per manufactures instructions. For ZC3H4 and WDR82 depletions, a mix of four siGENOME siRNAs was used (Catalog IDs MQ-066747-00 and MQ-062271-01, Horizon Discovery). As a control, the siGENOME Non-Targeting Control siRNA #2 (Horizon Discovery) was used. Cells were harvested 47 hours post transfection.

### Generation and validation of ZC3H4-dTAG cells

ZC3H4 was C-terminally tagged with a FKBP^12F36V^-2xHA-HiBit tag using CRISPR/Cas9-mediated genome editing. 400,000 F121-9 hybrid mESCs were co-transfected with 0.9 µg pHV177 and 1.8 µg pHV176 (plasmids described in Supplementary Table 1) in a suspension transfection using Lipofectamine 2000 (Thermo Fisher). GFP-positive cells were sorted, plated at low density and individual colonies were picked to establish clonal cell lines. Clonal lines were first screened for integration using the HiBit assay (Nano-Glo HiBiT Lytic Detection System, Promega), measured on the GloMax Discover (Promega). Homozygous lines were identified by genotyping selected lines using PlatinumII Hot-Start Green PCR mix (Invitrogen) and primers indicated in Supplemental Table 1. To find the earliest time-point of near-complete ZC3H4 degradation, 14,000 cells were plated per well in a white 96-well plate and treated with 500 nM dTAG13 (Sigma-Aldrich) for the indicated times before performing the HiBit assay. The background signal detected in wells containing no-HiBit cells was subtracted, and the signal of dTAG13-treated cells was normalized to the DMSO-treated control of the same clonal cell line.

### Western Blotting

Cells were lysed in 1x Laemmli buffer + β-ME at 10,000 cells / µL, and lysates were run on 4-20% precast gels. After transfer to a nitrocellulose membrane and blocking, blots were incubated in primary antibody overnight, following by a 1-hour incubation with HRP-conjugated secondary antibody. Blots were incubated with SuperSignal West Pico PLUS Chemiluminescent Substrate (Thermo Fisher) and imaged on a ChemiDoc (MP) imaging system (BioRad). The following primary antibodies were used: ZC3H4 (1:700, HPA040934, Merck), WDR82 (1:1000, 99715, Cell Signaling), GAPDH (1:10,000, 10494-1-AP, ProteinTech), and Lamin B1 (1:5000, sc-374015, Santa Cruz).

### INSERT-seq library construction and data processing

#### INSERT-seq experiment

INSERT-seq was performed on a polyclonal pool of cells that contained a library of different insert sequences at the Pou5f1 uaRNA locus, where a reporter had been introduced, as described in Vlaming et al,. 2022. This library consisted of both sequences occurring in the mouse genome, and sequences that were generated *in silico* (either randomly or designed). The INSERT-seq experiment after knockdown (see above) was described as in Vlaming et al., 2022, with one additional step. A fixed amount of RNA spike-in, described below, was added to the cell suspensions in TRIzol. This allowed a quantitative comparison between the different conditions. Pooled libraries were sequenced paired-end using the Illumina NovaSeq platform. Libraries derived from genomic DNA obtained ∼8.5M sequencing reads per sample. RNA-derived libraries were sequenced more deeply, with ∼42M reads per sample, to account for larger variation in abundance between inserts.

#### Generation of RNA spike-in for INSERT-seq

For spike-in control, we used *in vitro* transcribed RNA that had the same sequence as RNA transcribed from the Oct4 uaRNA reporter locus, containing the same flanking regions used to amplify the sequencing library, but with a few unique inserts that were not present in the library. These unique inserts were cloned into plasmids (see Supplementary Table 1) and PCR amplified to create a linear fragment with a T7 promoter. 100ng amplicon was used as a template for *in vitro* transcription by T7 RNA Polymerase (New England Biolabs) for 2 hours at 37°C in a 20 µL reaction with 1 mM per rNTP, 5 mM DTT and 0.5 µL RNase Inhibitor, murine (New England Biolabs). The produced RNA was added in variable amounts to identical samples of a known numbers of reporter-containing cells in TRIzol, and RT-qPCR was used to calculate that these cells contain ∼15 copies of RNA that are expressed from the Oct4 uaRNA reporter locus and long enough for the RT primer to anneal. For the INSERT-seq experiments, 0.12 copies of the spike-in transcript were added per cell in TRIzol; since ∼40% of cells contained an insert, expressed at ∼15 copies per cell, this led to ∼2% of reads mapping to one of the spiked-in insert sequences.

#### INSERT-seq data analysis

Raw reads were processed to counted reads per insert as described before (Vlaming et al., 2022). From the RNA counts tables, abundances of all unspliced and spliced versions of an insert were added up for a total abundance per insert per sample. Using DESeq2, these values were normalized for both insert abundance in the genomic DNA and spike-in abundance for each condition. Specifically, for the four spike-in sequence, the fraction of reads per sample mapping to each of them were added up, then averaged across the three replicates per condition. This average abundance per condition was multiplied by the total counts per sample to calculate a spike-in count per sample. Genomic DNA counts were normalized by setting the total of mapped reads to 1, averaged across 9 samples and inserts with very low abundance in (<10^-6^) were removed. The gDNA abundances (unique value per row) were multiplied by the calculated spike-in counts (unique value per column), and then the whole table was divided by the mean of all these factors. This table was used by DESeq2 to normalize the raw RNA counts.

Finally, to convert to more meaningful numbers before plotting, values were normalized to the average of the *in-silico*-generated random controls in cells treated with the non-targeting siRNA, so that those had a value of 1.

Normalized RNA values of below 5*10^-4^ were set to that value to allow log transformations. For analyses of inserts derived from the genome, the following filtering steps were done. First, inserts for which results were not reproducible between replicates were filtered out, i.e. inserts for which the standard deviation of the data was not below the average of the data for either the siNT or siZC3H4 condition. Then, inserts that were very lowly expressed in all conditions (0.5 or lower across all nine samples in total) were also filtered out.

Inserts in the library belonged to several categories. For the scatter plots in Figure 1C and 1D only inserts from a few predominant categories were plotted, namely inserts with random sequences (n=906) and sequences present in the mouse genome just downstream of TSSs (n=8,213) or around mRNA 3’ cleavage sites (n=215).

Homer (Heinz et al. 2010) was used to identify enriched motifs in unchanged inserts containing TSS-proximal/-distal mRNA-derived sequences (|L2FC|<0.25, N = 1,090). Inserts with TSS-proximal/-distal mRNA-derived sequences with L2FC > 0.5 (N = 2,121) were used as the background set. The Homer search was done in the top strand only (-norevopp), and motifs of 6, 8 or 10 nucleotides were considered.

Finding 5’ SS motifs and calculating their strengths was done as described before in Vlaming et al., 2022, by running MOODS (Korhonen et al. 2017) with lenient cutoffs and calculating MaxEnt scores (Yeo and Burge 2004). Classification of 5’ SSs in this manuscript was done using the cutoffs described by (Almada et al. 2013). MaxEnt scores of 4.0 - 7.39 were considered weak, 7.39 - 8.77 medium and > 8.77 strong.

To analyze the effect of 10-nucleotide (5’ splice site) sequences in different backgrounds, first the median abundance in the control condition (non-targeting siRNA) was calculated for all inserts with a scrambled sequence in a given background. All other inserts were normalized to this control level of the background sequence. Then, for every 10-nucleotide sequence, the mean effect was calculated over all inserts containing this sequence. For the plot stringently discarding any inserts showing signs of splicing, inserts for which even a single read mapped to the spliced version (defined in Vlaming et al., 2022) in the siNT condition was discarded before averaging the effects of each insert across backgrounds. Only medium/strong 5’ SSs were considered in this analysis.

The effects of mutant versions of introns within the INSERT-seq library was determined as described previously (Vlaming et al. 2022). Efficient splicing of the original intron (>70% spliced) and mutants effectively disrupting splicing (<10% spliced) were determined using the siNT data.

### TT-seq library construction and data processing

To generate TT-seq libraries after ZC3H4 KD, F121-9 cells were transfected with siRNAs as described above. Before harvesting at 47 hours post transfection, cells were exposed to 500 µM 4sU (SigmaAldrich, T4509) in fresh media for 20 minutes. 9 million cells were resuspended in 1.5mL Trizol to be used for TT-seq. Cell suspensions were spiked with 5% 4sU-labeled Drosophila S2 cells (2-hour labeling) resuspended in TRIzol based. TT-seq was performed as described in Mimoso and Adelman, 2023, with a minor modification at the fragmentation step. 60 µg RNA was fragmented in a 80 µL final volume for 3 minutes at 94°C. 350ng enriched RNA was used for library construction. Library construction was performed with the Illumina TruSeq stranded total RNA kit with RiboZero rRNA depletion, as described before (Mimoso and Adelman 2023). Libraries were pooled and sequenced paired-end on the Illumina NovaSeq 6000 platform.

To generate TT-seq libraries after rapid depletion of ZC3H4, ZC3H4-dTAG cells were treated with 500 nM dTAG13 (Sigma-Aldrich) for 40 minutes total and 500 µM 4sU was added for the last 10 minutes. Cells were washed with PBS, quickly trypsinized, quenched with cold DMEM + 10% FBS, and immediately placed on ice. All spins were performed at 4°C unless otherwise noted. Cells were spun down for 4 minutes at 1000 RPM, resuspended in 10 mL cold PBS, and counted. 30,000 cells were allocated for the HiBIT assay to confirm depletion of ZC3H4 in dTAG treated conditions, as described above. Cells were re-spun at 1000 RPM for 4 minutes and resuspended in 2 mL TRIzol. Samples were spiked with 5% 4sU-labeled *Drosophila* S2 cells (2-hour labeling) resuspended in TRIzol based on cell count. TT-seq libraries were constructed as in Mimoso and Adelman, 2023 with the following modifications: 200 ng of enriched RNA was used for library construction with the Illumina TruSeq stranded total RNA kit with RiboZero rRNA depletion, following manufacturer instructions for degraded RNA. Libraries were pooled and sequenced paired-end on the Illumina NextSeq500 platform.

Both KD and dTAG TT-seq datasets were processed as follows: Using a custom script (trim_and_filter_PE.pl), FASTQ read pairs were trimmed to 50 bp per mate for KD samples and 42 bp per mate for dTAG samples. Read pairs with a minimum average base quality score of 20 retained. Read pairs were further trimmed using cutadapt 1.14 to remove adapter sequences and low-quality 3’ bases (--match-read-wildcards -m 20 -q 10). Reads were first mapped the dm6 version of the *Drosophila* genome using STAR 2.7.31. Reads not mapping to the spike genome were then used for alignment to mouse (mm10) using parameters --quantMode TranscriptomeSAM GeneCounts --outMultimapperOrder Random --outSAMattrIHstart 0 -- outFilterType BySJout --outFilterMismatchNmax 4 --alignSJoverhangMin 8 --outSAMstrandField intronMotif --outFilterIntronMotifs RemoveNoncanonicalUnannotated --alignIntronMin 20 -- alignIntronMax 1000000 --alignMatesGapMax 1000000 --outFilterScoreMinOverLread 0 -- outFilterMatchNminOverLread 0. Duplicates were also removed using STAR. Stranded coverage bedGraph files were generated from deduplicated BAM files using STAR.

BedGraphs were normalized using the normalize_bedGraph custom script. The following normalization factors were used to depth normalize the siNT and siZC3H4 TT-seq libraries:

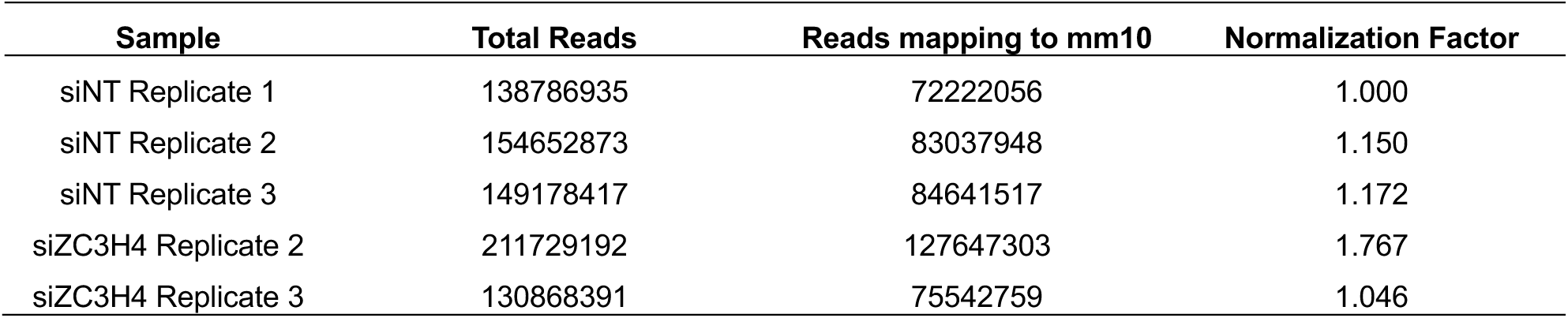

The following normalization factors were used to depth normalize the DMSO and dTAG TT-seq libraries:

| Sample | Number Total raw reads | Reads mapping to mm10 | Normalization Factor |
| --- | --- | --- | --- |
| DMSO Clone A6 | 61496524 | 58207811 | 1.205 |
| DMSO Clone A9 | 51240219 | 48972853 | 1.014 |
| DMSO Clone D7 | 56188487 | 53365028 | 1.105 |
| dTAG Clone A6 | 51003550 | 48286142 | 1.000 |
| dTAG Clone A9 | 67923594 | 64648250 | 1.339 |
| dTAG Clone D7 | 70774674 | 67300687 | 1.394 |

BedGraph files were converted to the bigWig format, and merged bedGraphs for each experimental condition were generated using bigWigMerge (UCSC tools). Merged bedGraphs were then converted to the bigWig format for visualization using bedGraphToBigWig (UCSC tools).

### PRO-seq library construction and data processing

Cells were permeabilized as described previously (Mimoso and Goldman 2023). Frozen (-80°C) permeabilized cells were thawed on ice, pipetted gently to fully resuspend, counted and aliquoted (Final concentration of 1 million cells per 45 µL). For each sample, 1 million permeabilized cells were used for nuclear run-on with 50,000 permeabilized *Drosophila* S2 cells added for normalization. Nuclear run on assays and library preparation were performed as described in Mimoso and Goldman, 2023. Pooled libraries were sequenced using the Illumina NextSeq platform.

All custom scripts described herein are available on the AdelmanLab GitHub (https://github.com/AdelmanLab/NIH_scripts). Dual, 6nt Unique Molecular Identifiers (UMIs) were extracted from read pairs using UMI-tools [10.1101/gr.209601.116]. Read pairs were trimmed using cutadapt 1.14 to remove adapter sequences (-O 1 --match-read-wildcards -m 26). The UMI length was trimmed off the end of both reads to prevent read-through into the mate’s UMI, which will happen for shorter fragments. An additional nucleotide was removed from the end of read 1 (R1), using seqtk trimfq (https://github.com/lh3/seqtk), to preserve a single mate orientation during alignment. The paired end reads were then mapped to a combined genome index, including both the spike (dm6) and primary (mm10) genomes, using bowtie2 [10.1038/nmeth.1923]. Properly paired reads were retained. These read pairs were then separated based on the genome (i.e. spike-in vs primary) to which they mapped, and both these spike and primary reads were independently deduplicated, again using UMI-tools. Reads mapping to the reference genome were separated according to whether they were R1 or R2, sorted via samtools 1.3.1 (-n), and subsequently converted to bedGraph format using a custom script (bowtie2stdBedGraph.pl). We note that this script counts each read once at the exact 3’ end of the nascent RNA. Because R1 in PRO-seq reveals the position of the RNA 3’ end, the “+” and “-“ strands were swapped to generate bedGraphs representing 3’ end positions at single nucleotide resolution.

BedGraphs were normalized using the normalize_bedGraph custom script. For libraries generated in DMSO- or dTAG- treated cells: We observed a consistent and significant decrease in reads mapping to the fly genome (dm6) after loss of ZC3H4 (Average ratio of dTAG / DMSO = 0.791; p-value = 0.011), indicating a global increase in transcription in dTAG-treated cells compared to the DMSO control. Accordingly, the number of reads mapping to the spike genome (dm6) was used to generate spike normalization factors, as shown below.

| Sample | Reads Mapping to dm6 | Reads Mapping to mm10 | Ratio of the % Spike Return (dTAG/DMSO) | Normalization Factor |
| --- | --- | --- | --- | --- |
| DMSO Clone A6 | 1248815 | 57107547 | 0.806 | 1.000 |
| dTAG Clone A6 | 1370661 | 64530686 |  | 1.098 |
| DMSO Clone A9 | 1883874 | 71288061 | 0.809 | 1.509 |
| dTAG Clone A9 | 1549062 | 73135770 |  | 1.240 |
| DMSO Clone D7 | 2036134 | 75094163 | 0.758 | 1.630 |
| dTAG Clone D7 | 1649135 | 83614278 |  | 1.321 |

Combined bedGraphs were generated by summing counts per nucleotide across replicates for each condition (bedgraphs2stdBedgraph). BedGraphs were converted to the bigWig format for visualization using bedGraphToBigWig (UCSC tools).

Processed and normalized PRO-seq bigwig files from Stein et al., 2022 (GSE200698) were downloaded for analyses after rapid depletion of INTS11 (DMSO and PROTAC; Figures 6 and 7). bigWigToBedGraph (UCSC tools) was used to convert bigwigs to bedGraphs.

### RNA-seq library construction and data processing

To generate RNA-seq libraries after ZC3H4 KD, F121-9 cells were transfected with siRNAs as described above. After 47 hours, 1M cells were resuspended in TRIzol. ERCC spike-in was added and RNA was extracted using DirectZol columns (Zymo Research). Genomic DNA contamination was removed using DNAse (Invitrogen), and RNA was cleaned up by adding TRIzol and extracting RNA using DirectZol columns again. All samples had RIN scores of >9. 800 ng of purified RNA was used as input to the Illumina TruSeq stranded total RNA kit with Ribo-Zero Gold rRNA depletion. Library construction was performed with the Illumina TruSeq stranded total RNA kit with RiboZero rRNA depletion, as described before (Mimoso and Adelman, 2023). Libraries were pooled and sequenced paired-end 51 bp on the Illumina NovaSeq platform.

RNA-seq library processing was performed as described above for TT-seq with the following modifications: FASTQ read pairs were trimmed to 50 bp per mate. BedGraphs were depth normalized using the normalize_bedGraph custom script and the following normalization factors:

| Sample | Number Total raw reads | Reads mapping to mm10 | Normalization Factor |
| --- | --- | --- | --- |
| siNT Replicate 1 | 89992062 | 53345775 | 1.356 |
| siNT Replicate 2 | 70708354 | 51873550 | 1.319 |
| siNT Replicate 3 | 63338645 | 51075231 | 1.298 |
| siZC3H4 Replicate 1 | 56248008 | 39342069 | 1.000 |
| siZC3H4 Replicate 2 | 84954927 | 67544740 | 1.717 |
| siZC3H4 Replicate 3 | 67996502 | 50972358 | 1.296 |

BedGraph files were converted to the bigWig format, and merged bedGraphs for each experimental condition were generated using bigWigMerge (UCSC tools). Merged bedGraphs were then converted to the bigWig format for visualization using bedGraphToBigWig (UCSC tools).

### ChIP-seq data processing

FASTQs corresponding to ZC3H4 ChIP-seq libraries in mESCs were downloaded from GSE199801. Using a custom script (trim_and_filter_PE.pl), FASTQ read pairs were trimmed to 36bp per mate, and read pairs with a minimum average base quality score of 20 were retained. Reads were then mapped to the mouse genome (mm10) using bowtie version 1.2.2 (-v2 -k1 -- allow-contain -X1000 -p 5 –best). The custom script extract_fragments.pl was used to retain reads fragments corresponding to insert sizes of 50-500bp, remove duplicate reads and generate bedGraphs (25nt bins) reporting the read fragment center. The custom script bedgraphs2stdBedGraph was used to merge replicate bedGraphs (n=3). The merged bedGraph were then converted to the bigWig format for visualization using bedGraphToBigWig (UCSC tools).

H3K27Ac ChIP-seq signal used in Figure S4 was processed as described in Vlaming et al., 2022.

### PAC-seq data processing

PAC-seq bedGraphs and bigWigs in exosome depleted cells (siRRP40) were generated as described in Mimoso and Adelman, 2023 (GSE218134).

### Gene and Enhancer Annotation

An annotation of the single dominant transcription start site (TSS) and transcription end site (TES) per active gene in mESCs was obtained as described in Mimoso and Adelman 2023 with the following modifications: PRO-seq data from DMSO and dTAG conditions and RNA-seq from WT mESCs was used as input for the custom script GetGeneAnnotation (available on the Adelman Lab Github (get_gene_annotations.sh, https://github.com/AdelmanLab/GeneAnnotationScripts). RNA-seq data from WT mESCs was used to avoid any unanticipated defects in 3’-end processing after loss of ZC3H4 from impacting isoform counts. Furthermore, uaRNA TSSs that overlapped an annotated TSS were removed from downstream analysis. Final number of genes and uaRNAs investigated are listed in the corresponding figure legends.

An annotation of active enhancers was generated as described in Stein et al., 2022 with the following modifications: Unnormalized PRO-seq bedGraphs from DMSO and dTAG-treated conditions were merged per strand. The merged bedGraphs were converted to bigwig files, and used as input for dREG (Danko et al. 2015). dRIP-filter (custom script; DOI 10.5281/zenodo.6654472) was run to filter the dREG output with the following parameters (-s 0.5 -p 0.025 -c 10). Bedtools intersect was used to remove dREG peaks within 1 kb of a dominant TSS (described above) or an annotated TSS in the basic GTF. Next, dREG peaks overlapping rRNA, snRNA, scRNA, srpRNA, and tRNA annotations in RepeatMaker were removed using bedtools intersect. Next, bedtools intersect was used to flag dREG peaks that overlapped an active gene. dREG peaks overlapping an active gene were flagged as an intragenic enhancer. dREG peaks that did not overlap an active gene were classified as an intergenic enhancer. Next, the dominant TSS was called within intergenic and intragenic dREG peaks. First, bedtools intersect was run to determine the overlap between dREG peaks and data-derived TSSs positions (“Annotated_Dominant_and_Nondominant_obsTSS_fordREG.txt” output file from GetGeneAnnotation). For intergenic dREG peaks, TSSs identified on both strands were considered when defining the dominant TSS position. For intragenic dREG peaks, only TSSs found on the strand opposite of the overlapping gene were considered when defining the dominant TSS position. This filtering step was applied to intragenic dREG peaks to distinguish between effects of ZC3H4 on the intragenic enhancer and the active gene. Next, only dominant TSSs classified as an unannotated TSS (nuTSS) by TSScall were retained. Lastly, ATAC-seq reads in mESCs (Processed data files from (Martin et al. 2023); GSE198517, 0 hour mESCs) were counted between 500 nt upstream of the dominant TSS to 500 nt downstream of the dominant TSS (TSS +/- 500 nt; 1 kb window). Enhancers with at least 50 ATAC-seq reads were retained. This generated a final working list of n=12,255 enhancers with a single dominant TSS (Intergenic = 9,548; Intragenic = 2,707).

### Box plots and Statistical Analysis

GraphPad Prism was used to generate box plots. Unless otherwise noted, all box plots have a line at the median and whiskers that depict 1.5 times the interquartile range. GraphPad Prism was also used to calculate P values, test used for each comparison is indicated in the figure legend.

### Scatter density plots

The get_density R function (http://slowkow.com/notes/ggplot2-color-by-density) and ggplot2 were used to generate density scatter plots in R (3.6.2).

### 5’ SS Strength (MaxEnt)

Score5.pl in the Maxentscan suite (https://github.com/Congenica/maxentscan.git) was used to calculate the strength of first 5’ SSs of intron-containing mRNAs (Figure 2E) relative to the consensus 5’ SS motif. Score5.pl was run as described in Mimoso and Adelman, 2023.

### CpG islands

The coordinates of CpG islands in mESCs was downloaded from UCSC table browser. Bedtools intersect was used to determine the overlap between CpG islands and the endogenous genomic locations of the sequences investigated by INSERT-seq. Number of inserts per species that overlap a CpG island is listed in the corresponding figure legends.

### Differential expression analysis

For mRNAs and lncRNAs, TT-seq reads within exons were summed per gene using featurecounts. For uaRNAs, TT-seq reads were summed between the uaTSS to +3 kb downstream. For eRNAs, TT-seq reads were summed between the eTSS to +2 kb downstream. Next, DESeq2 was used to generate a list of differentially expressed genes after loss of ZC3H4 (KD or dTAG). DESeq2 size factors were overwritten to match depth normalization. For calling differentially expressed genes between dTAG and DMSO conditions, design= ∼ batch + condition was applied, where batch corresponded to the different ZC3H4-dTAG clonal lines (A6, A9, D7). Affected genes were those with a log2 Fold Change > 0.50 and padj < 0.01. Genes with a log2 fold change < |0.25| were defined as unchanged. The number of affected genes per biotype are annotated in the corresponding figure legends.

### Heatmaps and metagene plots

The custom script make_heatmap (https://doi.org/10.5281/zenodo.5519915) was used to generate count matrices aligned to TSSs (mRNA TSS, uaTSS, lncRNA TSS or eTSS). Heatmaps were visualized using the Partek Genomics Suite. For all heatmaps, the bin size, number of visualized annotations and parameter used to rank the investigated data is indicated in the figure legends. Metagene plots were generated by summing reads within bins at each indicated position with respect to the TSS and dividing by the number of annotations. GraphPad Prism was used to visualize metagene plots. For all metagene plots, the bin size and number of investigated annotations are indicated in the figure legends.

### Conservation Score and Gene Age

The mm10 placental mammals base wise conversion by PhyloP dataset from UCSC Genome Browser was used to calculate the conservation scores of upregulated and unchanged mRNA TSSs in Figure 3F. A bigwig file was downloaded from UCSC table browser. bigWigToBedGraph (UCSCtools) was used to convert the PhyloP dataset bigWig file to a bedGraph. Gene ages used in Figure 3G were downloaded from (Zhang et al. 2010)(Data from Supplemental Table 2).

### Gene Downstream of uaRNA/eRNA

Bedtools closest was used to pair the closest active gene downstream of a uaTSS or eTSS (-iu -D a). ncRNAs with an mRNA gene within 5kb downstream of the uaTSS or eTSS were retained. Next, data was separated based on the directionality of the uaRNA or eRNA with respect to the downstream mRNA. Lastly, the fold change in TT-seq between dTAG- and DMSO-treated cells (as generated in Figure 3C) is reported for mRNAs downstream of upregulated uaTSS or eTSSs.

### K-means Clustering

Gene body PRO-seq reads were summed after rapid depletion of ZC3H4 (DMSO- and dTAG-treated cells, this study) or INTS11 (DMSO- and PROTAC-treated cells; Stein et al., 2022). mRNAs with at least 5 gene body PRO-seq reads in all 4 conditions and a log2FC > 0.5 in gene body PRO-seq signal after loss of ZC3H4 or INTS11 were retained. Next, gene body PRO-seq reads were normalized within each dataset (ZC3H4 or INTS11) such that the condition with the highest counts per gene was set to a maximum value of 1. The kmeans function in R (3.6.2) was used to define four clusters. For Figure 6A, genes within each cluster were ordered at random.

### Elongation Rate

Elongation rate was calculated by dividing TT-seq read coverage (RNA synthesis) by PRO-seq 3’ end reads (the signal representing RNAPII density), as described in Mimoso and Adelman 2023. A constant factor of 1 was added to each heatmap to avoid dividing by 0.

### Heatmaps and metagene plots

The custom script make_heatmap (https://doi.org/10.5281/zenodo.5519915) was used to generate count matrices aligned to TSSs (mRNA TSS, uaTSS, lncRNA TSS or eTSS). Heatmaps were visualized using the Partek Genomics Suite. For all heatmaps, the bin size, number of visualized annotations and parameter used to rank the investigated data is indicated in the figure legends. Metagene plots were generated by summing reads within bins at each indicated position with respect to the TSS and dividing by the number of annotations. GraphPad Prism was used to visualize metagene plots. For all metagene plots, the bin size and number of investigated annotations are indicated in the figure legends.

### Conservation Score and Gene Age

The mm10 placental mammals base wise conversion by PhyloP dataset from UCSC Genome Browser was used to calculate the conservation scores of upregulated and unchanged mRNA TSSs in Figure 3F. A bigwig file was downloaded from UCSC table browser. bigWigToBedGraph (UCSCtools) was used to convert the PhyloP dataset bigWig file to a bedGraph. Gene ages used in Figure 3G were downloaded from (Zhang et al. 2010)(Data from Supplemental Table 2).

### Gene Downstream of uaRNA/eRNA

Bedtools closest was used to pair the closest active gene downstream of a uaTSS or eTSS (-iu-D a). ncRNAs with an mRNA gene within 5kb downstream of the uaTSS or eTSS were retained. Next, data was separated based on the directionality of the uaRNA or eRNA with respect to the downstream mRNA. Lastly, the fold change in TT-seq between dTAG- and DMSO-treated cells (as generated in Figure 3C) is reported for mRNAs downstream of upregulated uaTSS or eTSSs.

### K-means Clustering

Gene body PRO-seq reads were summed after rapid depletion of ZC3H4 (DMSO- and dTAG-treated cells, this study) or INTS11 (DMSO- and PROTAC-treated cells; Stein et al., 2022). mRNAs with at least 5 gene body PRO-seq reads in all 4 conditions and a log2FC > 0.5 in gene body PRO-seq signal after loss of ZC3H4 or INTS11 were retained. Next, gene body PRO-seq reads were normalized within each dataset (ZC3H4 or INTS11) such that the condition with the highest counts per gene was set to a maximum value of 1. The kmeans function in R (3.6.2) was used to define four clusters. For Figure 6A, genes within each cluster were ordered at random.

### Elongation Rate

Elongation rate was calculated by dividing TT-seq read coverage (RNA synthesis) by PRO-seq 3’ end reads (the signal representing RNAPII density), as described in Mimoso and Adelman 2023. A constant factor of 1 was added to each heatmap to avoid dividing by 0.

---

**Supplemental Figure S1.**
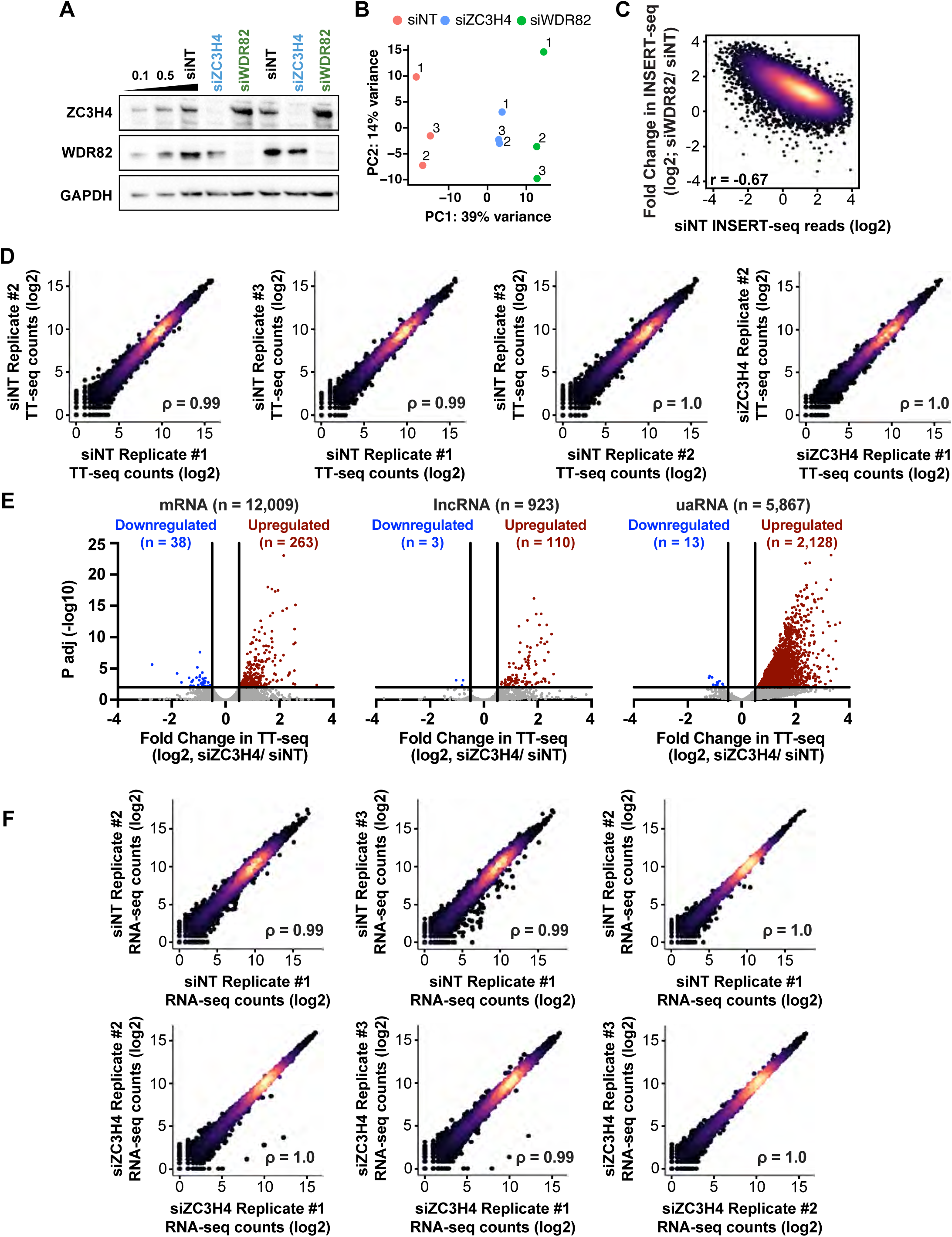
Depletion of Restrictor increases synthesis and abundance of many ncRNAs. Related to Figure 1. *(A)* mESCs were transfected with siRNAs targeting Restrictor subunits ZC3H4 or WDR82 or a non-targeting control (NT) for 48 hrs. Western blots show the protein levels of ZC3H4, WDR82 and GAPDH in the corresponding conditions. GAPDH is shown as a loading control. *(B)* PCA plot reporting the agreement between INSERT-seq replicates. For this plot, the fully normalized insert abundances (see methods) of all inserts analyzed in this manuscript were used. *(C)* Density scatter plots reporting the fold change in INSERT-seq levels between siWDR82 and siNT conditions graphed against INSERT-seq expression levels under siNT conditions (n = 9,334). *(D)* Density scatter plots reporting the replicate agreement between TT-seq samples generated after siZC3H4. TT-seq reads were counted over exons. Data is shown for active genes (n = 13,088). *(E)* Volcano plot per gene biotype depicting differentially expressed genes in siZC3H4 and siNT treated cells. TT-seq reads were calculated within exons for mRNAs and lncRNAs. For uaRNAs, TT-seq reads were counted between the uaTSS to 3 kb downstream. Affected genes were defined by DESeq2 (p <0.01 and Fold Change (log2) >0.50). *(F)* Same as D, except for RNA-seq samples generated after transfection with siZC3H4 or siNT (n = 13,088).

**Supplemental Figure S2.**
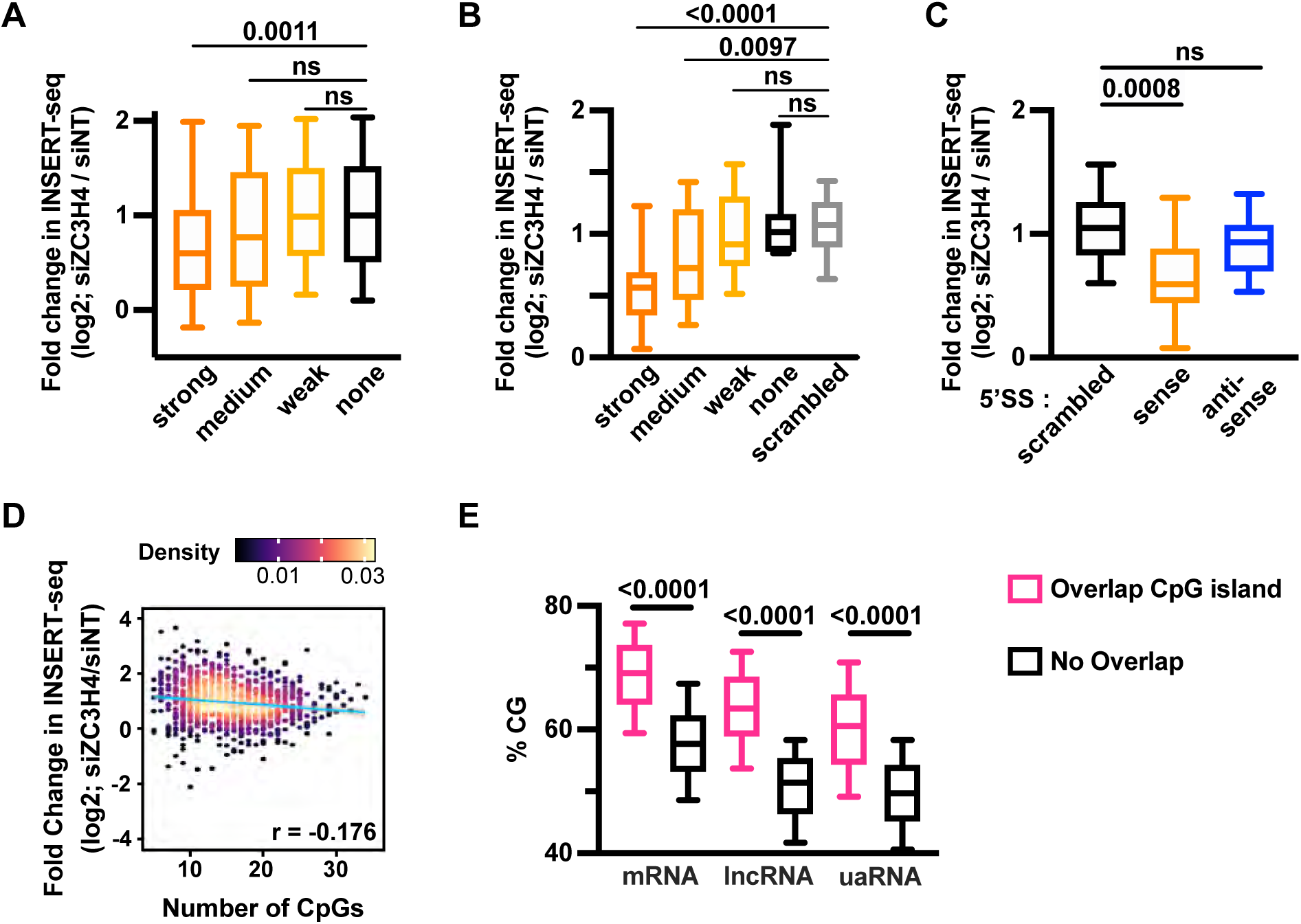
5’ Splice sites, rather than CpG islands or CG content, govern sensitivity to Restrictor. Related to Figure 2. *(A)* Box plot reporting the fold change in INSERT-seq signal between siZC3H4 and siNT conditions for synthetic sequences grouped by the presence and strength (MaxEnt score) of 5’ SS motif matches they contained by chance (none, n = 398; weak, n = 338; medium, n = 106; strong, n = 64). Box plots have a line at the median with whiskers from 10-90th percentile. Mann-Whitney test used to generate p-values. ns = not significant, p > 0.05. *(B)* As in A, but showing synthetic sequences with annotated 5’ SS sequences inserted, and each data point is based on the average effect of the same splice donor sequence in up to five different backgrounds. Inserts were separated based on the strength (MaxEnt score) of inserted 5’ SS motifs and compared to sequences with a scrambled 5’ SS sequence inserted (scrambled, n = 30; none, n = 7; weak, n = 15; medium, n = 19; strong, n = 19). *(C)* Same as in Fig. 2C, but filtering out inserts for which any spliced reads could be detected before averaging the results per 5’ SS across backgrounds (scrambled, n = 18; sense, n = 36; antisense, n = 18). *(D)* Density scatter plot reporting the fold change in INSERT-seq after ZC3H4 KD with respect to number of CpGs per insert. Data is shown for all synthetic sequences evaluated here (n = 906). *(F)* Box plots reporting the % CG content for groups shown in Fig. 2G, with TSS-proximal inserts from each RNA biotype separated based on overlap with a CpG island. Data is shown per biotype (left to right, n = 2918, 691, 129, 179, 819, 588). Mann-Whitney test used to generate p-values.

**Supplemental Figure S3.**
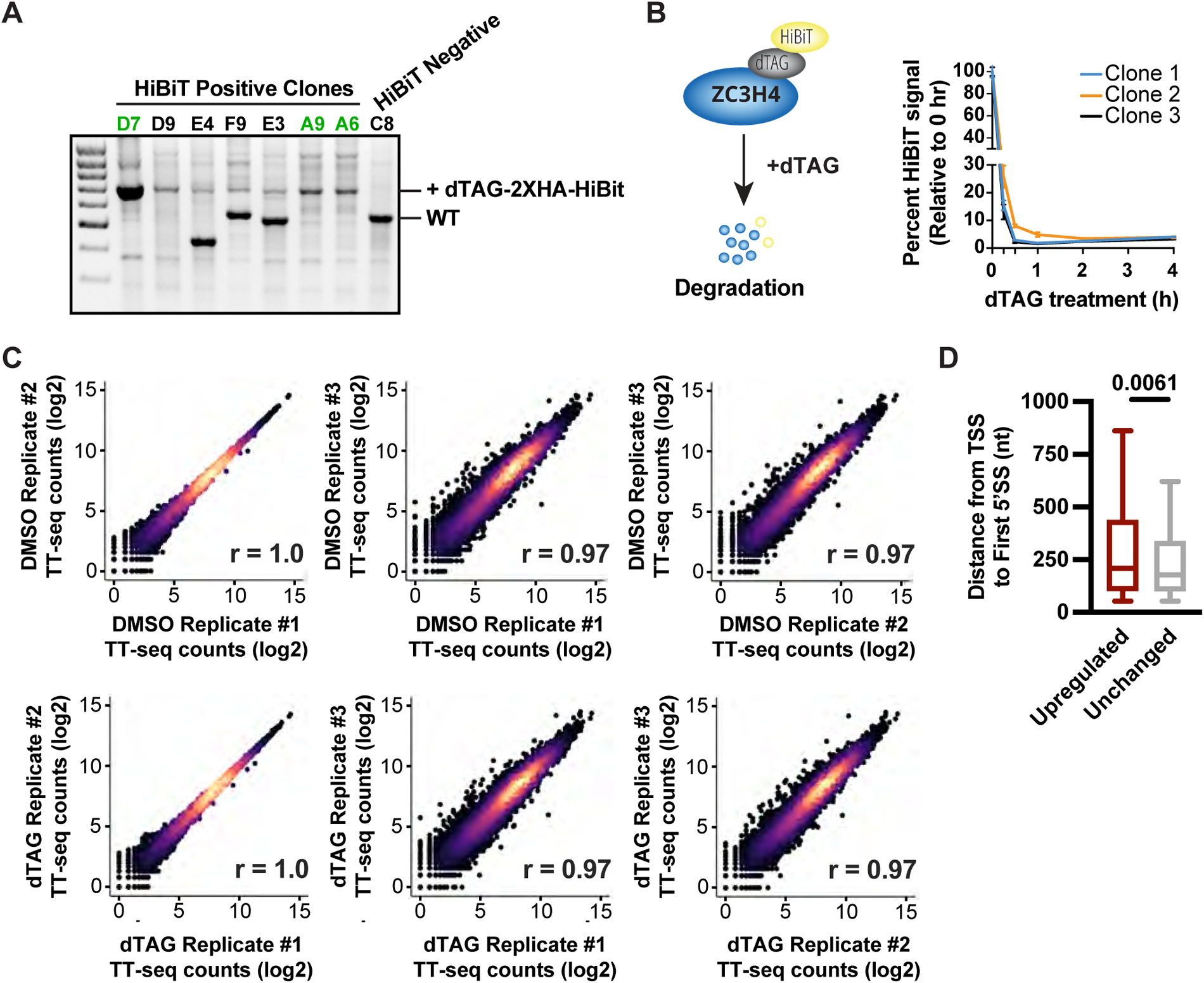
Rapid degradation of ZC3H4 followed by analyses of newly synthesized RNA supports a co-transcriptional role for Restrictor. Related to Figure 3. *(A)* HiBiT signal was used to identify cell clones with successful integration. Genotyping PCR is shown for HiBiT positive and negative clones. The size of PCR fragments corresponding to WT ZC3H4 and ZC3H4-dTAG-2XHA-HiBit is indicated. The three clones with homozygous integration of the dTAG-2xHA-HiBIT used in this study are highlighted in green. *(B)* ZC3H4 abundance was measured using the HiBiT fluorescence assay. Signal for three independent clones is shown, normalized to the 0-hr time point. Error bars indicate SD (n=3). *(C)* Density scatter plots reporting the replicate agreement between TT-seq samples generated after dTAG-13 or DMSO treatment (n = 3 per condition). TT-seq reads were counted over exons. Data is shown for all active genes (n = 13,088). *(D)* Box plots depicting the distribution of distances between the TSS and first 5’ SS at upregulated (n = 441) and unchanged (n= 7,547) mRNAs. Box plots have a line at the median with whiskers from 10-90th percentile. P-values from the Mann-Whitney test.

**Supplemental Figure S4.**
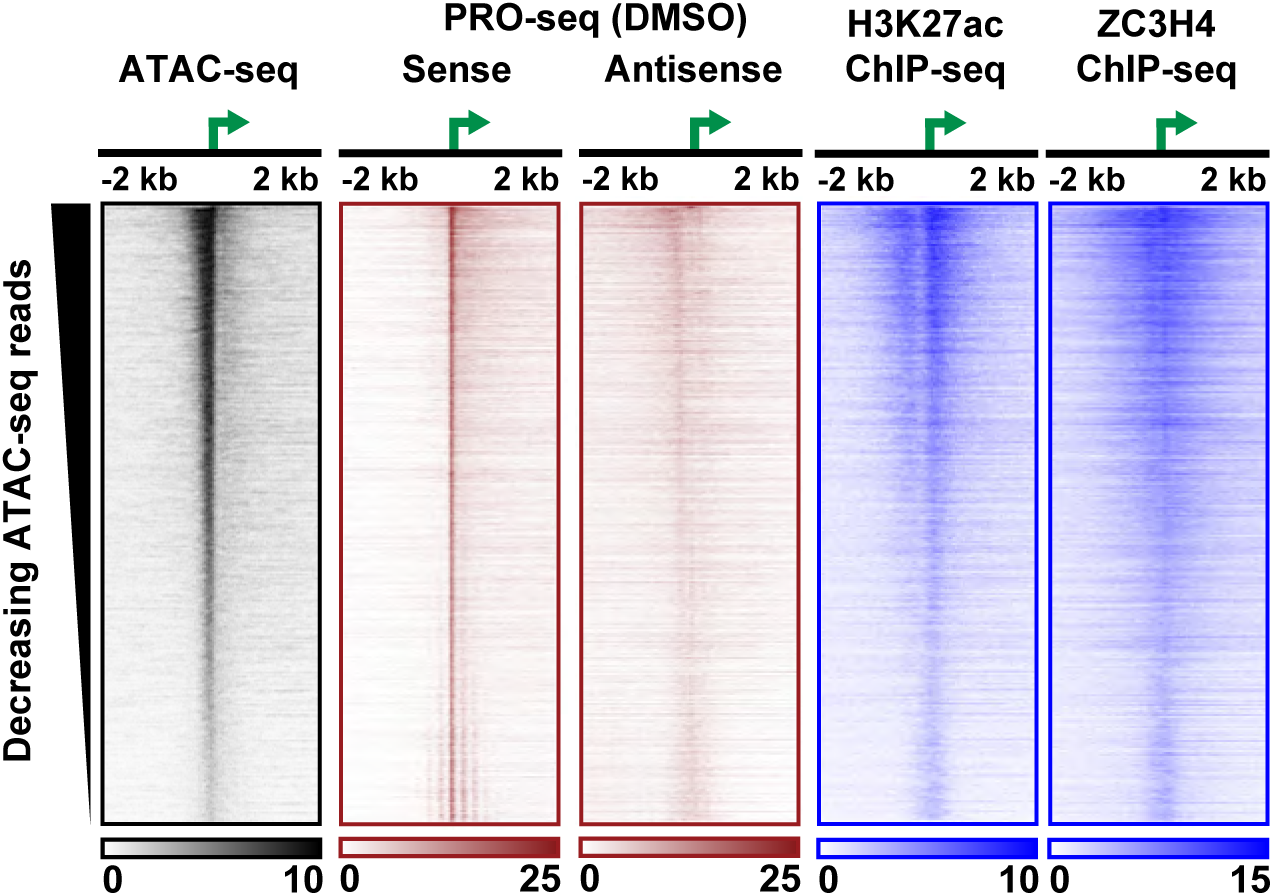
Identification of enhancers in mESCs and confirmation that ZC3H4 localizes to these regions. Heatmaps depicting the indicated datasets at active enhancers, defined using ATAC-seq from control mESCs and PRO-seq from DMSO- and dTAG-treated cells (n = 12,255). Data is aligned to the dominant TSS within each enhancer (called here the eTSS), as defined by 5’ ends of PRO-seq reads. ATAC-seq reads were counted from eTSS +/- 500 bp. eRNAs are ranked by decreasing number of ATAC-seq reads in this window. For the heatmaps, ATAC-seq and PRO-seq were summed in 25 nt bins. ChIP-seq for H3K27 acetylation (from Vlaming et al. 2022) and ZC3H4 (from Hughes et al. 2023) was summed in 50 nt bins.

**Supplemental Figure S5.**
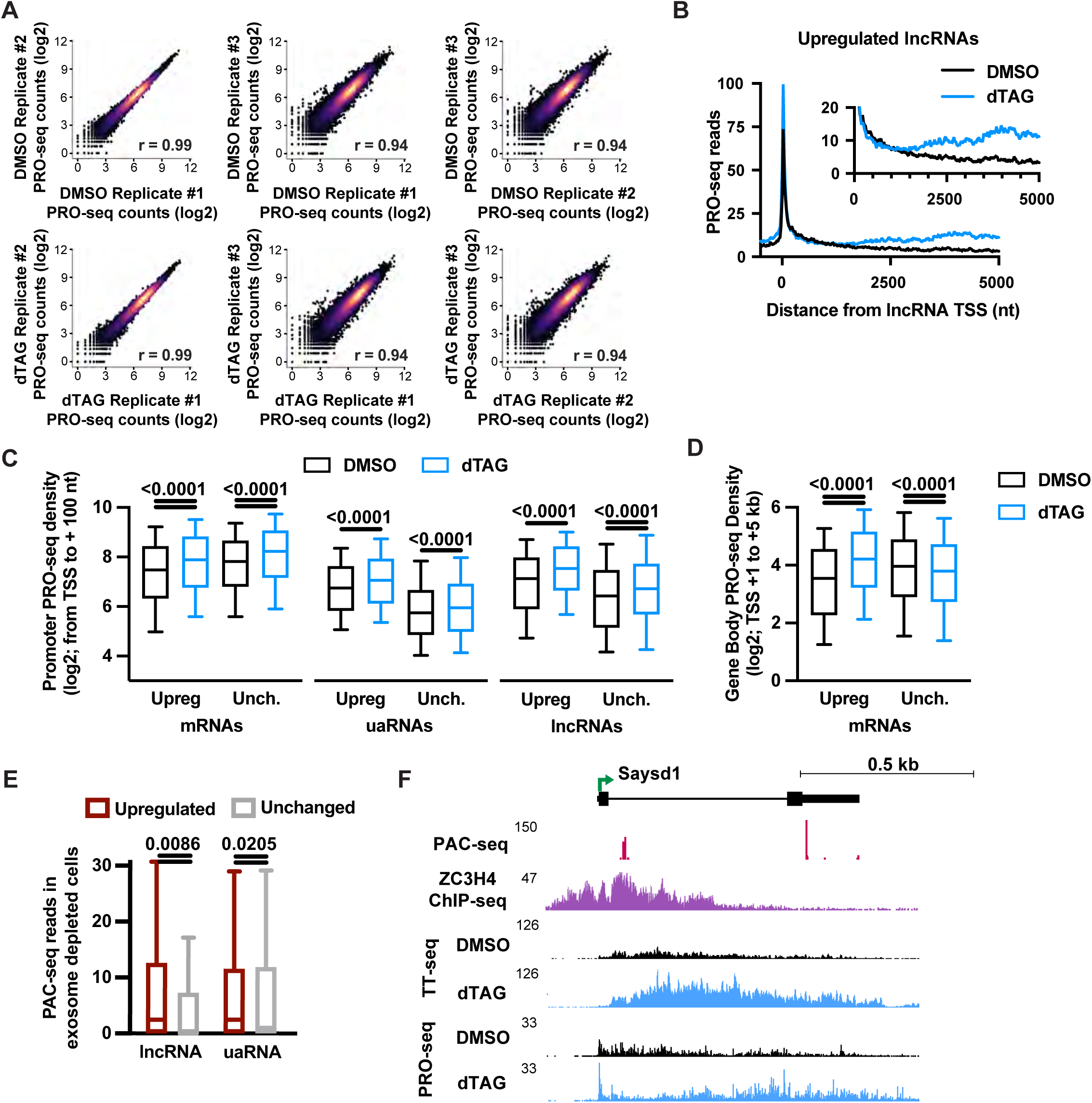
Acute depletion of Restrictor causes locus-specific changes in RNAPII elongation. Related to Figure 5. *(A)* Density scatter plots reporting replicate agreement between PRO-seq samples generated after dTAG- or DMSO-treatment (n = 3 per condition). PRO-seq reads were counted between the TSS and 100-nt downstream. Data is shown for active genes (n = 13,088). *(B)* Metagene plots of PRO-seq signal in dTAG- and DMSO-treated cells at upregulated lncRNAs. Data is aligned to the TSS, and summed in 25-nt bins. Inset highlights gene body PRO-seq signal. *(C)* Box plots reporting promoter PRO-seq density (TSS to +100-nt) at the indicated gene lists. Box plots have a line at the median with whiskers from 10-90th percentile. P-values were generated using the Wilcoxon test. *(D)* Box plots reporting gene body PRO-seq density (from +1 kb to +5 kb downstream of the indicated mRNA TSSs). P-values were generated using the Wilcoxon test. We note that in contrast to the upregulated genes, the unchanged genes (defined by TT-seq) have a modest reduction in PRO-seq signal in this window. *(E)* Box plots report PAC-seq reads in cells depleted of the exosome (using siRRP40) to inhibit RNA decay. Reads were summed between TSS and + 1250-nt downstream at upregulated and unchanged lncRNAs and uaRNAs. Box plots have a line at the median and whiskers depicting 1.5 times the interquartile range. P-values were generated using the Mann-Whitney test. *(F)* Example mRNA (Saysd1) upregulated after loss of ZC3H4. Data is shown on the sense strand for indicated data types and conditions.

**Supplemental Figure S6.**
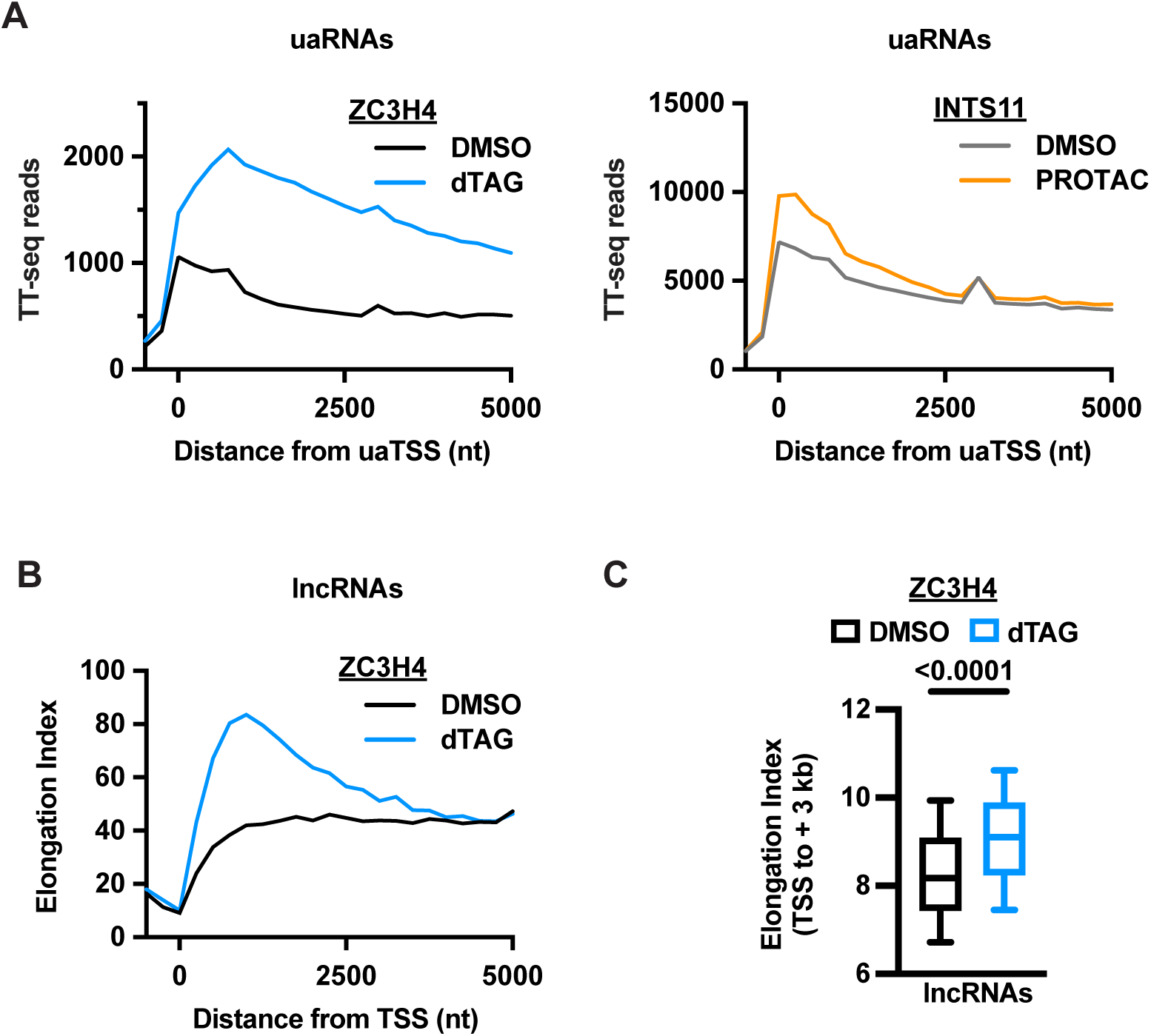
Restrictor reduces RNAPII elongation rate. Related to Figure 7. *(A) Left:* Metagene plots of TT-seq signal in ZC3H4-tagged cells after DMSO or dTAG treatment, at uaRNAs (n = 5,867). Data is aligned to the uaTSS, and summed in 250-nt bins. *Right:* Same as left, except showing INTS11-tagged cells after DMSO or PROTAC treatment. *(B)* Metagene plots report Elongation Index in DMSO- and dTAG-treated ZC3H4-dTAG cells at lncRNAs (n = 923). Elongation Index was calculated as the ratio of TT-seq signal (RNA synthesis) to PRO-seq signal (elongating RNAPII) per 250-nt bin. *(C)* Box plot reporting the sum of Elongation Index values between TSS to +3 kb downstream in DMSO- or dTAG-treated ZC3H4-dTAG cells at lncRNAs (n = 923). Box plots have line at median with whiskers from 10-90th percentile.

**Supplementary table 1:**
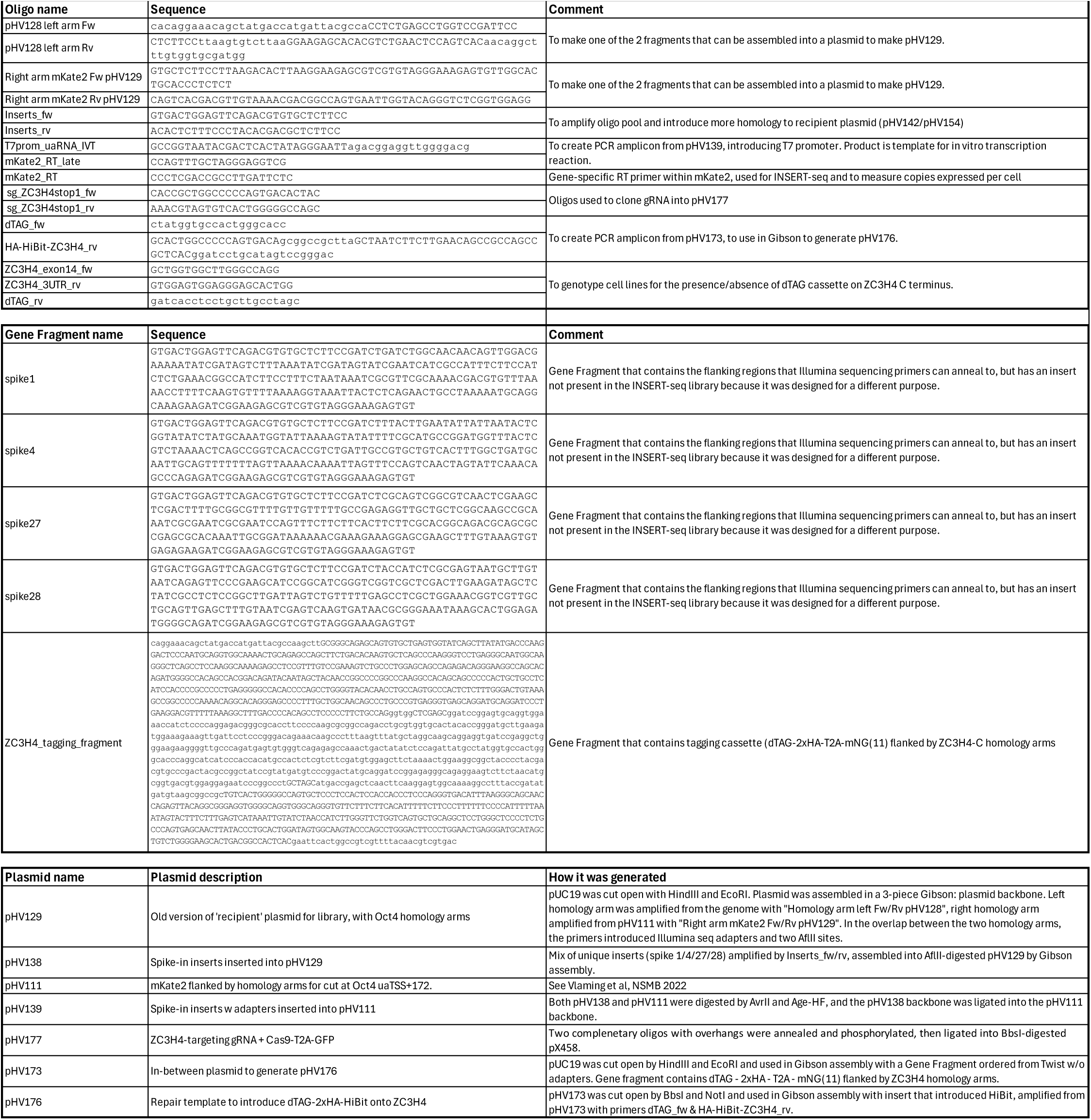
oligos, gene fragments and plasmids.

